# Microglial SWELL1 Deficiency Drives Male-Specific Seizure Vulnerability but Paradoxical Neuroprotection through Impaired Phagocytosis

**DOI:** 10.1101/2025.05.26.656163

**Authors:** Abhijeet S. Barath, Aastha Dheer, Emily Dale, Flavia Goche, Thanh Thanh Le Nguyen, Laura Montier, Mastura Akter, Mekenzie Peshoff, FangFang Qi, Anthony D. Umpierre, Dale B. Bosco, Koichiro Haruwaka, Rajan Sah, Long-Jun Wu

## Abstract

The discovery of genes encoding the volume-regulated anion channel (VRAC) has enabled detailed exploration of its cell type–specific roles in the brain. LRRC8A (SWELL1) is the essential VRAC subunit. We observed seizure-induced, subunit-specific changes in microglial VRAC expression and investigated its function using conditional knockout (cKO) of LRRC8A in microglia. SWELL1 cKO mice exhibited a male-specific increase in kainate-induced seizure severity yet showed paradoxical neuroprotection against seizure-associated neuronal loss. Mechanistically, SWELL1 deletion led to a cell-autonomous reduction in microglial density and decreased release of VRAC-permeable neuroactive metabolites, including taurine, GABA, and glutamate in culture. Additionally, impaired phagocytic kinetics and reduced lysosomal biogenesis contributed to the observed neuroprotection. These findings reveal novel roles for microglial VRAC in regulating seizure outcomes and microglia–neuron interactions.

## INTRODUCTION

Microglia, as the primary immune cells of the central nervous system, have long been studied for their role in neuroinflammation and neurodegeneration following seizures (*1–3*). The emergence of seizures as a clinical manifestation in primary microgliopathies such as Nasu-Hakola disease (linked to TREM2/DAP12 dysfunction) and adult-onset leukoencephalopathy with axonal spheroids and pigmented glia (ALSP; associated with CSF1R haploinsufficiency) has further underscored their relevance in acute seizure pathophysiology (*4, 5*). Interest in this area has intensified with findings showing that microglial ablation exacerbates kainate-induced seizures in adult mice (*6–8*). Several microglial signaling pathways, including P2Y12R-ATP, CX3CR1-CX3CL1, TREM2, and Gi-coupled GPCRs, have been implicated in modulating seizure severity (*1, 2, 7, 9–14*). Notably, recent clinical data has shown that P2Y12 receptor polymorphisms influence seizure phenotype in patients, lending translational significance to these preclinical findings (*15*).

Following a seizure, microglia contribute to tissue remodeling by clearing damaged neurons via phagocytic receptors and releasing inflammatory cytokines (*3, 11, 16*). Microglial TREM2 and P2Y6, as well as Gi coupled GPCR signaling have been directly implicated in regulating neuronal loss after seizures (*11, 12, 16*). Whether microglial phagocytosis is beneficial or detrimental remains debated and likely depends on the outcome measured. For instance, P2Y6 deficiency impaired microglial phagocytosis, resulting in reduced neuronal loss which was associated with improved hippocampal-dependent memory in the weeks following status epilepticus (*16*). However, TREM2 loss, which was also associated with impaired phagocytosis and neuroprotection, led to an increased risk of spontaneous recurrent seizures (SRS) (*11*). Similarly, microglia play a role in promoting aberrant neurogenesis following seizures, which increases susceptibility to SRS (*2, 3, 11, 17–22*).

The volume regulated anion channel (VRAC), is a heterogenous, multi subunit channel which classically mediates regulatory volume decrease of cells under conditions of hyposmotic stress (*23*). It is encoded by the members of Lrrc8 gene family (Lrrc8a-e) (*24, 25*). The SWELL1 protein, encoded by the *Lrrc8a* gene, is essential for functional VRAC channel formation (*23–25*). In our previous study, bulk RNA-sequencing following kainate-induced status epilepticus revealed complex changes in the expression of all subunits of VRAC expressed in microglia (*11*). VRAC is broadly expressed across mammalian cell types and recent studies have highlighted its diverse and context-dependent roles, including regulating astrocytic glutamate release, influencing stroke outcomes, and mediating mechanosensation in vascular endothelium (*24–28*). Pharmacological studies have further implicated VRAC in cellular migration, motility, and proliferation (*29–31*)—functions particularly relevant to microglial responses during seizures. These findings prompted us to investigate the role of microglial VRAC in acute seizures and subsequent pathology. Here, we employ advanced transgenic mouse models to selectively delete *Lrrc8a* in microglia. Our results demonstrate that microglial SWELL1 critically modulates both acute seizure severity and post-seizure neuropathology likely via mechanistically distinct pathways.

## RESULTS

### Seizures downregulate microglial SWELL1 expression while microglia-specific SWELL1 deficiency leads to increased seizure severity

Microglia are known to regulate neuronal excitability during acute seizures and coordinate neuroimmune responses during the postictal period (*1–3, 21*). To explore the role of microglial VRAC in this context, we analyzed our previously published bulk RNA sequencing data from CD11b⁺ cells isolated from wild-type mice before and 7 days after kainate (KA)-induced seizures (*11*). This analysis revealed significant alterations in homeostatic and seizure-associated gene expression, as well as in all Lrrc8 family members expressed in microglia (Fig. 1A). At baseline, Lrrc8a and Lrrc8d exhibited the highest expression levels among the VRAC subunits, with lower levels of Lrrc8b and Lrrc8c (Fig. 1B). A similar subunit expression pattern was observed at the protein level in ex vivo and primary cultured microglia using publicly available proteomic data (Fig. S1C) (*32*). Seven days following seizure induction, Lrrc8a (Swell1), the obligate VRAC subunit, was significantly downregulated, while Lrrc8 subunits b–d were upregulated (Fig. 1B), suggesting seizure-induced reconfiguration of VRAC subunit composition in microglia.

**Figure 1.**
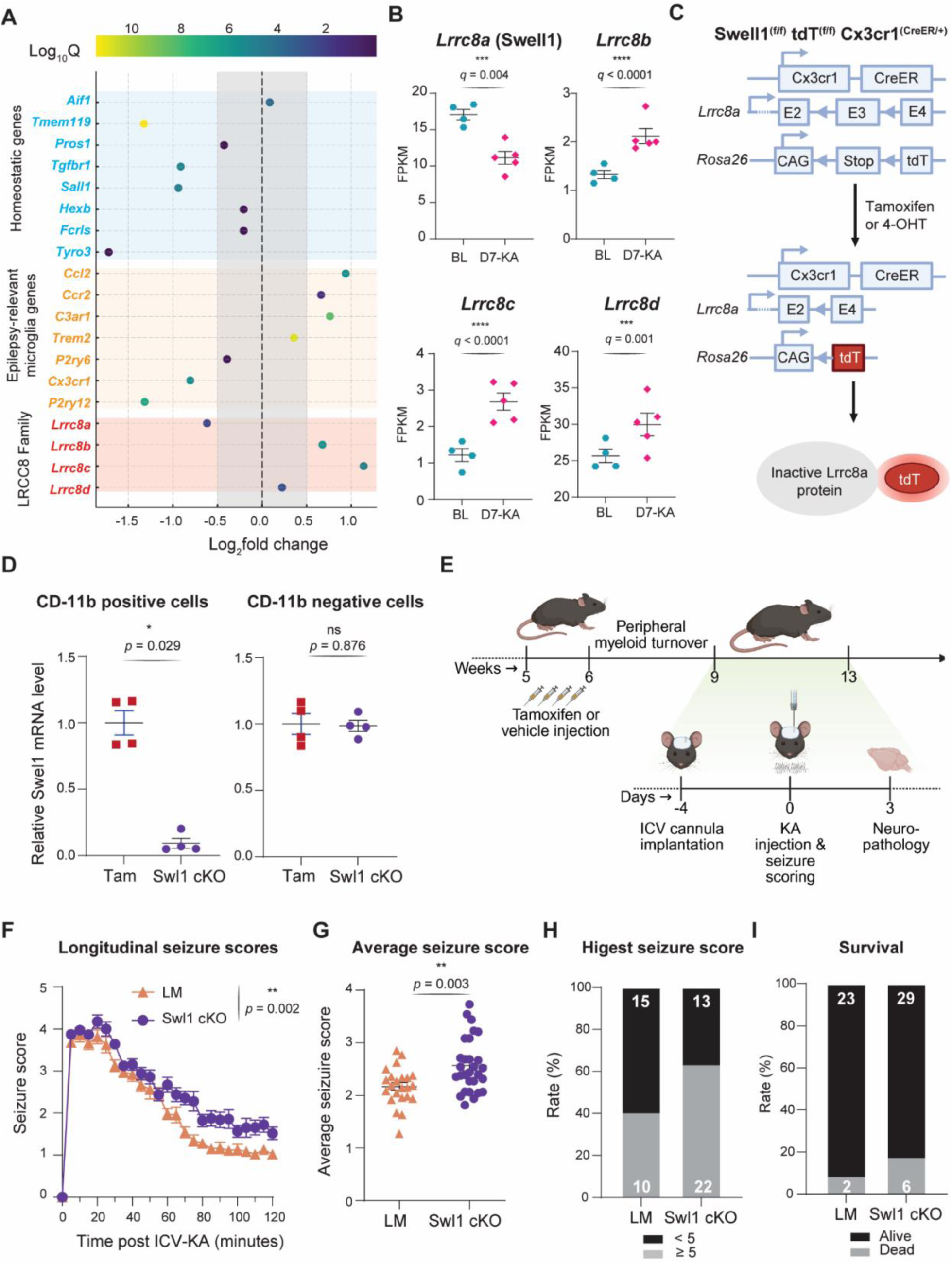
Bidirectional interplay between microglial VRAC expression and seizure severity. A) A dot plot showing changes in microglial homeostatic, epilepsy related, and LRRC8 family of genes at day 7 after kainate induced seizure in wild type mice (bulk-sequencing of brain CD11b^+^ cells; n = 4-5 WT mice/group) B) LRRC8a (Swell1) – LRRC8d expression in microglia at day 7 after seizures (statistic: q value i.e., false discovery rate (FDR) adjusted p-value) C) Genetics of Swell1 conditional knockout mice-stop codons are placed around the E3 exon which is spliced out after tamoxifen or 4-OHT treatment. tdTomato expression is also activated D) Relative Swell1 mRNA levels in CD11b^+^ and CD11b^-^ cells showing cKO efficiency and selectivity (mRNA normalized to GAPDH), vs tamoxifen (tam) treated control animals (statistic: Mann-Whitney test and unpaired t-test) E) Seizure experiment timeline F) Longitudinal Racine seizure scores in Swell1 cKO vs littermate (LM) controls (n=25-35 mice/group; statistic: 2-way ANOVA) G) Average seizure scores over the two-hour observation period (statistic: unpaired t-test) H) Percent of mice achieving a Racine score of 5 or higher at least once. I) Survival rate. 4-OHT: 4-hydroxy tamoxifen (active metabolite of tamoxifen); ICV: intracerebroventricular; KA: kainate; LM: vehicle treated littermate controls; Tam: Tamoxifen treated controls.

To investigate the functional consequences of this shift, we generated a tamoxifen-inducible, SWELL1 conditional knockout (cKO) by crossing SWELL1^(fl/fl)^ mice with CX3CR1^(CreER/CreER)^ mice (Fig. 1, C and E). This approach enabled selective deletion of SWELL1 in adult microglia. Quantitative PCR confirmed a robust and selective reduction in SWELL1 mRNA in CD11b⁺ brain cells following tamoxifen administration, achieving an average knockdown efficiency of 91 ± 8% (Fig. 1D). Functional *in vitro* validation demonstrated impaired regulatory volume decrease in SWELL1-deficient microglia when exposed to a mild hypotonic challenge (osmolarity 200-215 mOsm/L; Fig. S1, A and B; Vid. S1 and S2). Swell1 cKO microglia displayed much slower kinetics of volume reduction and larger cell sizes than controls.

To assess the in vivo impact of microglial SWELL1 loss, we employed the intracerebroventricular kainate (ICV-KA) model of temporal lobe epilepsy. Interestingly, we found male SWELL1 cKO mice exhibited significantly increased seizure severity compared to vehicle-treated littermate controls, as assessed by the modified Racine scale (Fig. 1, F and G). There were also non-significant trends toward higher rates of severe seizures (Racine score ≥5) and mortality in the cKO group (Fig. 1, H and I). These findings were replicated using tamoxifen-treated controls lacking either the Cre recombinase or floxed SWELL1 alleles (i.e., SWELL1^(fl/fl)^ CX3CR1^(wt/wt)^ and SWELL1^(wt/wt)^ CX3CR1^(CreER/wt)^), thereby ruling out off-target effects of tamoxifen (Fig. S1, D–G). Together, these results demonstrate that seizures alter microglial VRAC expression and abrogating VRAC expression increases severity of acute seizures in mice, suggesting a bidirectional relationship between seizure activity and microglial VRAC function.

### Transient microglial loss and morphological remodeling in SWELL1 cKO mice

Previous studies have implicated VRAC/SWELL1 in cell proliferation and survival (*29–31*). Consistent with these roles, we observed a ∼10–20% reduction in microglial density in both the cortex and the CA3 pyramidal layer of SWELL1 cKO mice at 4–6 weeks following tamoxifen induction (Fig. 2, A–D; Fig. S2, A and B). This reduction was accompanied by increased nearest neighbor distances (NND), indicative of altered microglial tiling, and a rightward shift in NND distribution curves (Fig. S2E). Importantly, microglial density returned to control levels by 10 weeks, confirming the transient nature of this reduction (Fig. 2, A and C). These findings were further validated by CreER-driven tdTomato expression (Fig. S2, C and D) and reduced Iba1 immunoreactivity at 4–6 weeks (Fig. 2E). At the time point of reduced microglial density, we also noted changes in microglial morphology. Specifically, microglial territory size was increased (Fig. 2F), and Sholl analysis revealed complex alterations in arborization patterns (Fig. 2G), suggesting structural remodeling in response to SWELL1 loss.

**Figure 2.**
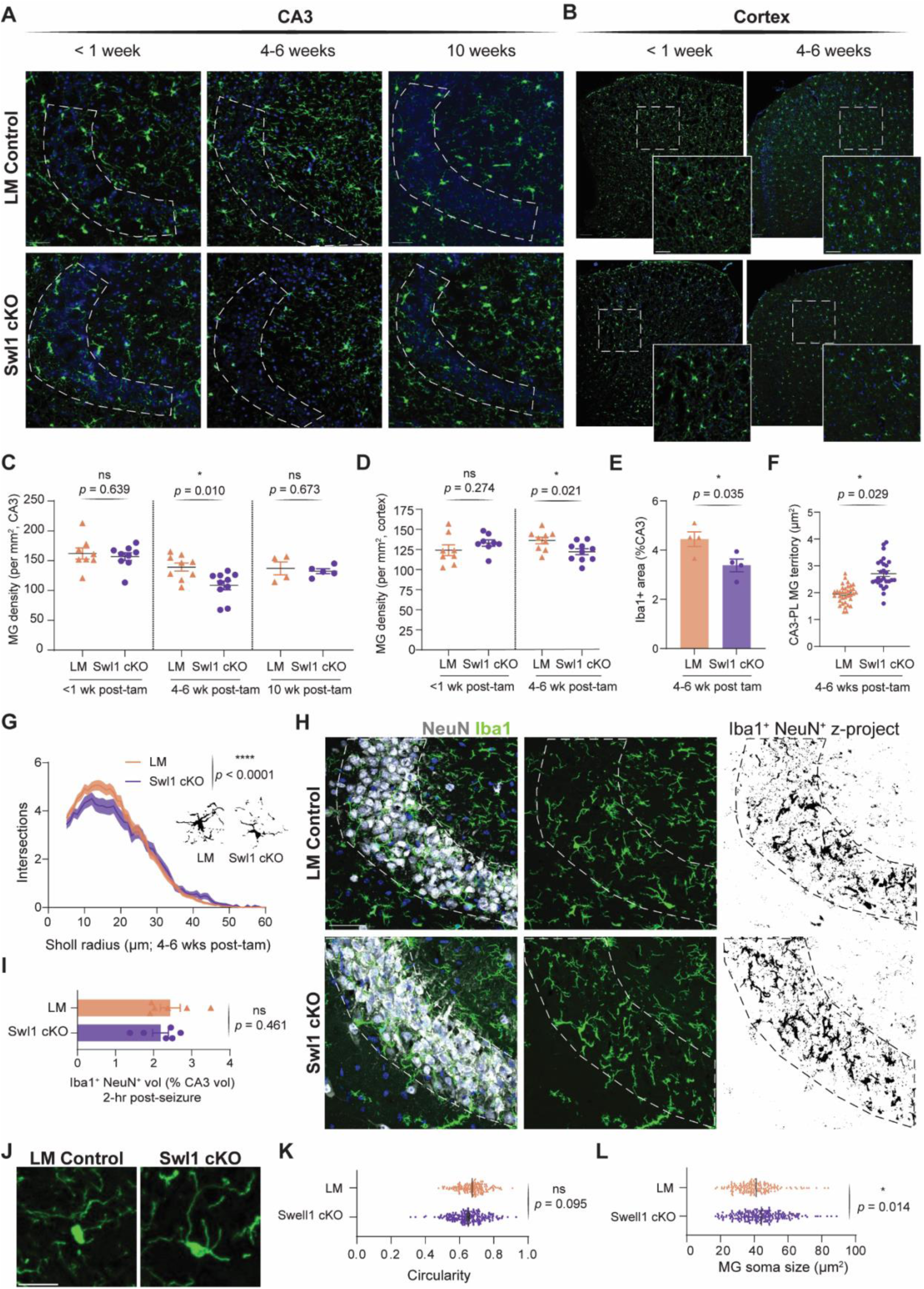
Transient microglial loss and morphological remodeling in SWELL1 cKO mice. A) Microglia density in CA3 pyramidal layer of hippocampus at various time points after the last tamoxifen/ vehicle administration. Scale bar, 50 µm. B) Microglia density in cortex at various time points after the last tamoxifen/ vehicle administration. Scale bar, 50 µm. C) Quantification of microglia density in CA3 pyramidal layer (statistic: unpaired t-test or Mann-Whitney test) D) Quantification of microglia density in cortex (statistic: unpaired t-test). E) Quantification of Iba1^+^ area in CA3 pyramidal layer (statistic: unpaired t-test). F) Quantification of microglia territory in CA3 pyramidal layer (dot, individual microglia; 10-40 microglia/mouse, 4-5 mice/group; statistic: nested t-test). G) Sholl analysis and representative thresholded images of microglia from Swell1 cKO and LM control mice (10-40 microglia/mouse, 4-5 mice/group; statistic: 2-way ANOVA). H) Microglia (Iba1) - neuron (NeuN) interaction in CA3 pyramidal layer at 2 hours after kainate induced seizure. Scale bar, 50 µm. I) Iba1-NeuN double positive volume as a percent of CA3 pyramidal layer volume (statistic: unpaired t-test) J) Microglia soma changes at 2-hour after seizure induction. Scale bar. 20 µm. K) Quantification of microglia soma circularity in CA3 (dot, individual microglia; 90-160 microglia/mouse, 5-6 mice/group; statistic: nested t-test). L) Quantification of microglia soma size in CA3 (dot, individual microglia; 90-160 microglia/mouse, 5-6 mice/group; statistic: nested t-test). LM: vehicle treated littermate control; PL: pyramidal layer; Tam: tamoxifen; wk: weeks;

Because these density and morphological changes coincided with the timing of our seizure behavioral experiments, we hypothesized that reduced microglial–neuronal contact might contribute to heightened seizure severity (Fig. S2F). To test this, we quantified microglia–neuron interactions two hours after acute seizures by calculating the percentage of CA3 voxels co-labeled with microglial markers (Iba1 or P2Y12) and the neuronal marker NeuN using high-resolution confocal microscopy (40x objective; voxel size: 0.21 × 0.21 × 0.5 µm³). No significant differences were observed between SWELL1 cKO and control mice (Fig. 2, H and I; Fig. S2, G and H). However, morphometric analysis revealed larger microglial soma size and a trend toward reduced soma circularity in SWELL1 cKO mice (Fig. 2, J–L), suggestive of altered activation states. Notably, soma sizes were reduced from baseline in both genotypes (Fig. S2, I and J). These morphological changes may be attributed to volume redistribution as microglia extend their processes and form bulbous endings or to a volume loss, raising the possibility that swelling-associated signaling mechanisms could contribute to the seizure phenotype in SWELL1-deficient mice.

### SWELL1 deletion impairs microglial release of neuroactive metabolites

VRAC facilitates regulatory volume decrease through the efflux of a variety of organic and inorganic anions, several of which possess neuromodulatory properties (*23, 24, 31, 33*). Thus, we hypothesized that impaired release of inhibitory metabolites from seizure-activated SWELL1 cKO microglia could contribute to the increased seizure severity observed in this genotype (Fig. S3C). To test this, we used a mild hypotonic stimulus to activate VRAC in cultured microglia and measured the release of 14 neuroactive metabolites (via LC-MS; Fig. 3A) and ATP (via colorimetric assay; Table S1). Eight metabolites, along with ATP, were reliably detected. Among these, the release of taurine, GABA, and glutamate was significantly reduced (∼50%) in SWELL1 cKO microglia under hypotonic conditions (Fig. 3, B–D), whereas the release of aspartate, adenosine, and ATP remained unaffected (Fig. 3, D–G).

**Figure 3.**
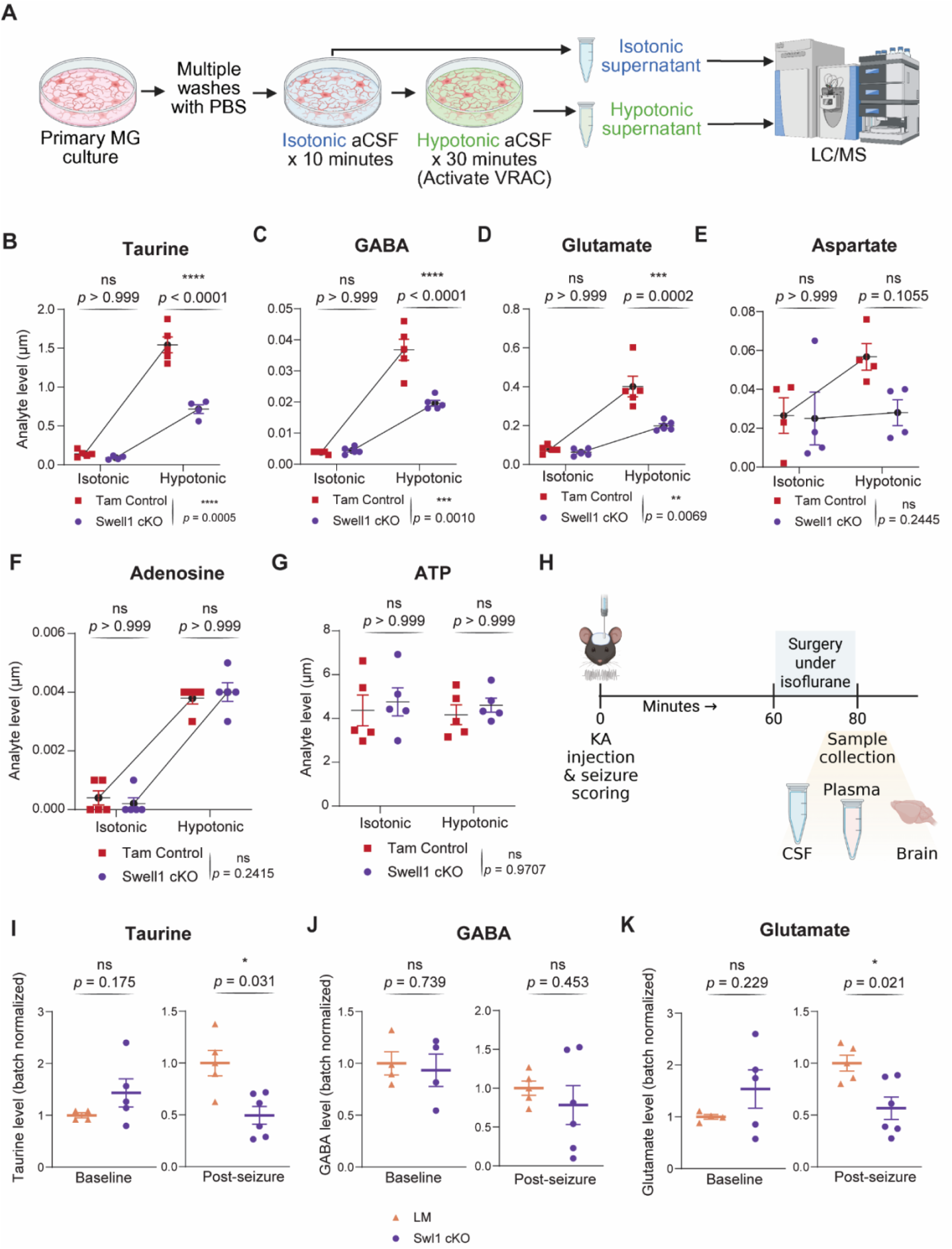
SWELL1 deletion impairs microglial release of neuroactive metabolites. A) Experiment design for activation of Swell1 (VRAC channel) in purified microglia cultures with a hypotonic stimulus and collection of supernatant for neuromodulator analysis. B) (B-G) quantification of taurine, GABA, glutamate, aspartate, adenosine, and ATP levels in supernatant of Swell1 cKO vs control microglia maintained in isotonic and hypotonic conditions (dot, one well; 4-5 wells/genotype; 50,000 microglia/well; statistic: 2-way Anova with Bonferroni’s multiple comparison test). H) Experiment design to test differences in CSF and plasma metabolites after seizure induction. I) (I-K) Batch normalized taurine, GABA, and glutamate levels (statistic: Permutation test). LC/MS: Liquid chromatography/Mass spectrometry; VRAC: volume regulated anion channel

To determine whether these effects are reflected *in vivo*, we analyzed cerebrospinal fluid (CSF) samples collected from mice at baseline and one hour after KA-induced seizures (Fig. 3H). Data were batch-normalized to minimize technical variability. At baseline, CSF levels of taurine, GABA, and glutamate were comparable between genotypes. However, following seizures, SWELL1 cKO mice exhibited significantly lower levels of taurine and glutamate compared to littermate controls (Fig. 3, I–K), while post-seizure GABA levels did not differ significantly (Fig. 3J). In both genotypes, post-seizure CSF levels of glutamate, GABA, and aspartate were strongly correlated (Fig. S3, D and E). Notably, taurine levels were highly correlated with these metabolites only in littermate controls, suggesting disrupted taurine regulation in SWELL1 cKO mice. Raw values for all CSF and plasma metabolite measurements are provided in Supplementary Tables S2–S5.

### SWELL1 deletion confers paradoxical neuroprotection following seizures

We next assessed the impact of SWELL1 cKO on seizure-induced neuronal loss. Using Nissl staining, unexpectedly, we found that SWELL1 cKO mice exhibited significantly reduced neuronal loss in the CA3 region of the hippocampus compared to littermate controls, as assessed 3 days after KA-induced seizures (Fig. 4, A and C). This neuroprotection was confirmed with NeuN staining (Fig. Bi, Biv, 4D). Importantly, baseline neuronal density did not differ between genotypes (Fig. S4, A–D). These findings were paradoxical, as SWELL1 cKO mice experienced more severe seizures.

**Figure 4:**
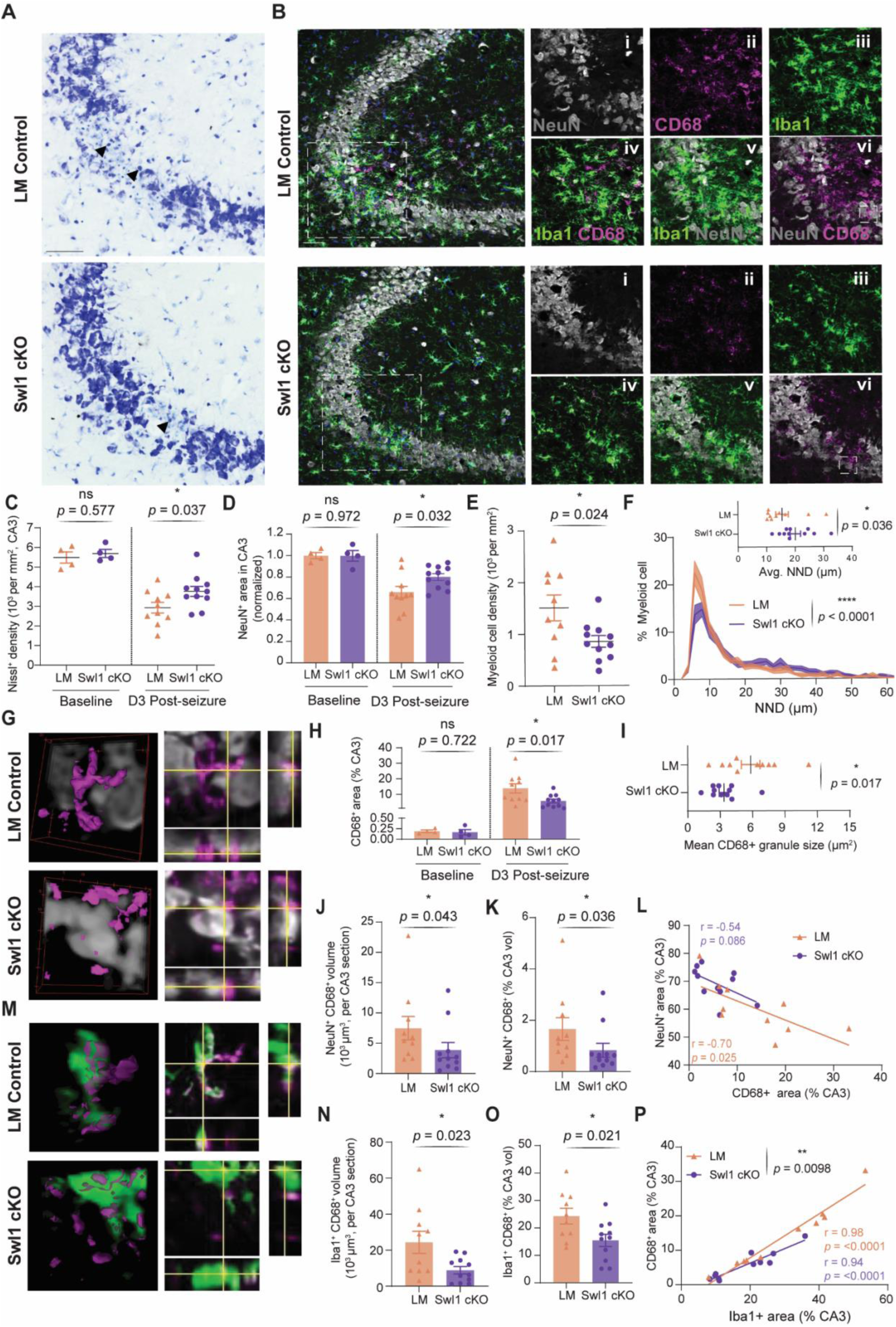
SWELL1 deletion confers paradoxical neuroprotection following seizures due to reduced lysosomal biogenesis. A) Nissl staining showing neuronal loss in CA3 pyramidal layer at day3 after seizure (arrowheads, pyknotic nuclei). Scale bar, 50 µm. B) Immunofluorescence images showing neuronal loss, myeloid and CD68 responses in CA3 at day 3 after seizure. Scale bar, 50 µm for images with overlay of all channels. C) Quantification of Nissl positive cell bodies at baseline and day 3 post seizure (statistic: unpaired t-test). D) Quantification of relative NeuN positive area in CA3 at baseline and day 3 after seizure (statistic: unpaired t-test). E) Quantification of myeloid cell density in CA3 pyramidal layer at day 3 after seizure (statistic: unpaired t-test). F) Distribution and average of nearest neighbor distances for myeloid cells in CA3 pyramidal layer (statistic: Mann-Whitney test and 2-way ANOVA). G) 3D reconstructed and orthogonal views of NeuN-CD68 volumetric interactions in CA3 pyramidal layer (zoomed in view of ROIs shown subpanel B.vi and B.xii). H) Quantification of CD68^+^ area in CA3 at baseline and day 3 after seizures (statistic: unpaired t-test). I) Quantification of mean CD68 granule sizes at day 3 after seizures. J) (J-K) Quantification of NeuN-CD68 double positive voxels in CA3 pyramidal layer (statistic: Mann-Whiteny test). L) Correlation between CD68 ^+^ and NeuN ^+^ area in CA3 pyramidal layer at day 3 after seizure (statistic: Pearson correlation coefficient) M) 3D reconstructed and orthogonal views of Iba1-CD68 volumetric interactions in CA3 pyramidal layer (zoomed in view of same area as the ROIs shown subpanel B.vi and B.xii). N) (N-O) Quantification of Iba1-CD68 double positive voxels in CA3 pyramidal layer (statistic: unpaired t-test) **P)** Correlation between Iba1^+^ and CD68 ^+^ area in CA3 pyramidal layer at day 3 after seizure (statistic: Pearson correlation coefficient (r); simple linear regression to compare the slopes)

Given that seizure-induced neurodegeneration is closely linked to microglial phagocytosis, we next examined the microglial and lysosomal responses. We found that Iba1⁺ cells aggregated in the CA3 region in both groups; however, SWELL1 cKO mice showed significantly lower microglial cell density and Iba1 immunoreactivity (Fig. 4, Biii, Bix, and E; Fig. S4, F and G). In line with lower cell density, nearest neighbor distances (NND) were significantly increased in SWELL1 cKO mice, with a rightward shift in the distribution (Fig. 4F), indicating more distant microglial tiling.

We next examined the lysosome marker CD68 in microglia. We found that SWELL1 cKO mice exhibited a blunted lysosomal response, as evidenced by significantly lower CD68 staining intensity and smaller, less numerous CD68⁺ granules in the CA3 region (Fig. 4, Bii, Bviii, G–I; Fig. S4H). Notably, the NeuN⁺ area in the CA3 pyramidal layer showed a strong inverse correlation with CD68 intensity in littermate controls (Fig. 4L), consistent with increased neuronal loss being associated with heightened microglial phagolysosomal activity. A moderate, non-significant negative correlation was observed in SWELL1 cKO mice, suggesting a disrupted link between neuronal loss and lysosomal engagement. To further investigate phagolysosomal function, we quantified the volume of double-positive NeuN⁺ CD68⁺ voxels—representing neuronal material in contact with or internalized by lysosomes—using high-resolution confocal imaging (20x magnification; voxel size: 0.43 × 0.43 × 1 µm³; Fig. 4G). SWELL1 cKO mice displayed a significantly smaller volume of NeuN⁺ CD68⁺ voxels compared to controls (Fig. 4, J and K), supporting reduced neuronal clearance as the basis for the observed neuroprotection.

As CD68 expression is closely tied to microglial activation and proliferation, we examined whether reduced CD68 staining in SWELL1 cKO mice could be explained by fewer microglia. We quantified the volume of Iba1⁺ CD68⁺ voxels as a proportion of total Iba1⁺ volume. Both the absolute and relative volume of these double-positive voxels were significantly reduced in SWELL1 cKO mice (Fig. 4, N and O). Moreover, while CD68⁺ and Iba1⁺ areas were positively correlated in both genotypes, the slope of this relationship was significantly lower in SWELL1 cKO mice (Fig. 4P), indicating an impairment in efficiency of lysosomal biogenesis. The data in Fig. 4L and 4P represent averages pooled from 3–5 CA3 sections per animal. Notably, substantial variability in Iba1 and CD68 responses was observed between sections from the same animal, suggesting regional heterogeneity in microglial activation. To account for this, we conducted a secondary analysis treating individual sections as biological replicates. This approach confirmed the strong correlations between CD68⁺ and NeuN⁺ areas (Fig. S4I), and between Iba1⁺ and CD68⁺ areas (Fig. S4J), including the reduced slope in SWELL1 cKO mice, thus reinforcing our conclusions regarding reduced efficiency of lysosomal synthesis in the latter.

### Cell-autonomous proliferative and phagocytic deficits in SWELL1 cKO microglia

To determine whether the proliferative and phagocytic impairments observed in SWELL1 cKO mice were cell-autonomous, we performed *in vitro* experiments using primary cultured microglia. When plated at equal density, SWELL1 cKO microglia exhibited ∼25% lower cell numbers than tamoxifen-treated controls by day 10 in culture (Fig. 5, A and B), indicating a proliferation defect. Because phagocytosis can be influenced by cell density, we performed the phagocytosis assay at an earlier time point, prior to the emergence of significant density differences (Fig. S5A). GFP⁺ latex beads opsonized with fetal bovine serum (FBS) IgG were used as the phagocytic substrate (Fig. 5C). Microglia were activated during the assay due to incubation in serum-free medium, which reduces Fcγ receptor saturation (normally caused by serum IgG), thereby enhancing sensitivity to IgG-opsonized beads (Fig. 5D). We found that SWELL1 cKO microglia displayed significantly impaired phagocytic kinetics compared to tamoxifen-treated controls, as measured by both the proportion of microglia containing bead inclusions and the average phagocytic load per cell (Fig. 5, D–F). These findings were independently confirmed using vehicle-treated, genotype-matched controls (Fig. S5, B–D). Together, these results demonstrate that the proliferative and phagocytic defects in SWELL1-deficient microglia are intrinsic and cell-autonomous.

**Figure 5.**
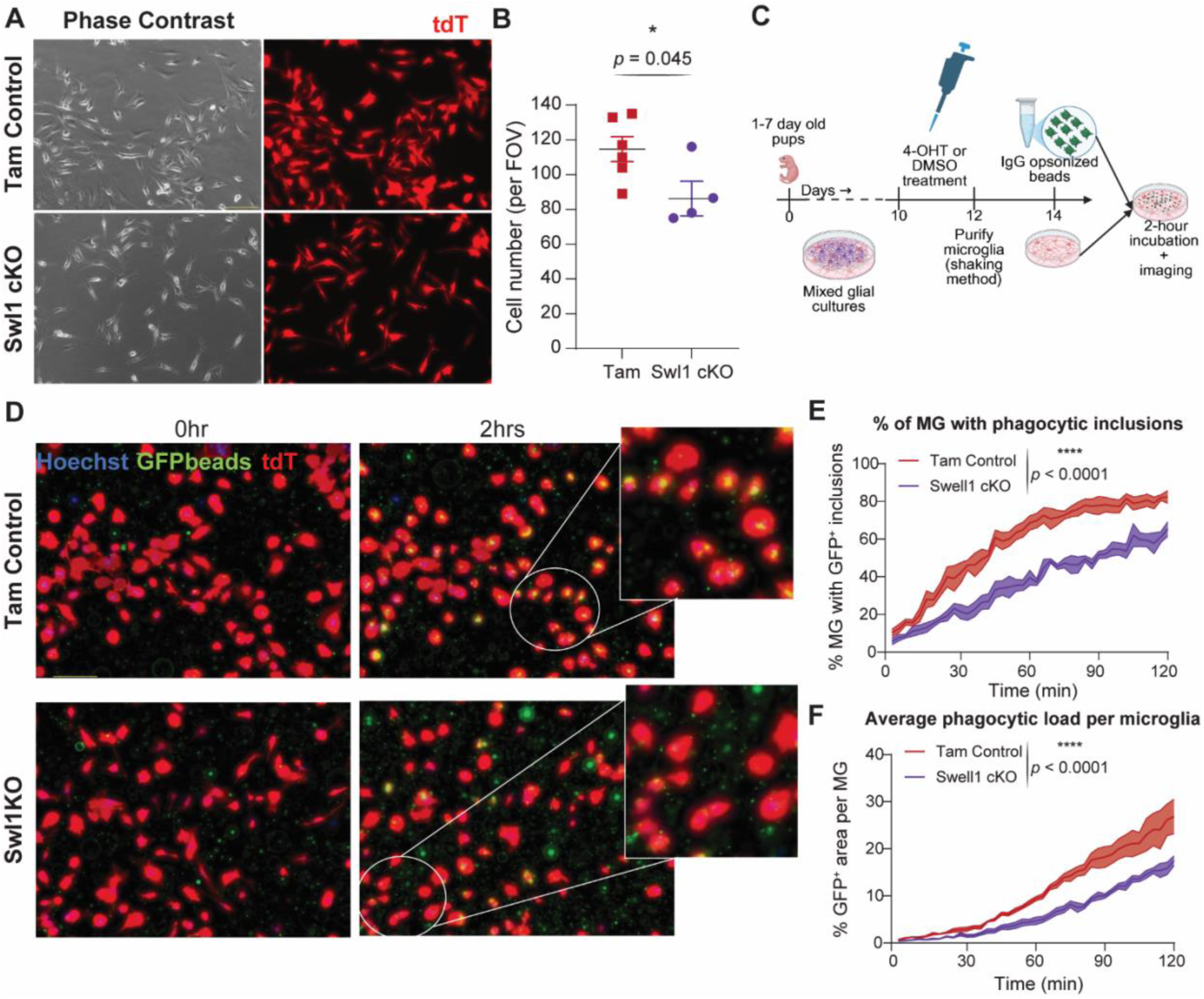
Cell-autonomous proliferative and phagocytic deficits in cultured SWELL1 cKO microglia. A) Microglia density at 10 days *in vitro* after plating purified microglia at equal density. Scale bar, 100 µm. B) Quantification of microglia density at 12 days *in vitro* (statistic: unpaired t-test). C) Experiment design for *in vitro* phagocytosis assay with opsonized latex beads. D) Bead phagocytosis assay. Scale bar, 100 µm. E) Quantification of percent microglia that are positive for phagocytic inclusions over a two-hour observation window (n = 4 wells/genotype; statistic: 2-way ANOVA) F) Quantification of average phagocytic load per microglia expressed as GFP ^+^ area (n = 4 wells/genotype; results averaged per well; statistic: 2-way ANOVA)

### Female SWELL1 cKO mice exhibit neuroprotection without increased seizure severity

Sex differences in microglial function have been reported in the contexts of neuropathic pain, phagocytosis, and P2Y12 dependent behaviors (*31, 34–36*). Therefore, we examined its role in Swell1 dependent effects. Interestingly, unlike male SWELL1 cKO mice, females showed no differences in seizure severity or mortality compared to littermate controls (Fig. 6, A and B; Fig. S6, A and B). Although this trend was noted during early experiments, all female data were analyzed separately to systematically highlight both similarities and sex-specific differences.

**Figure 6.**
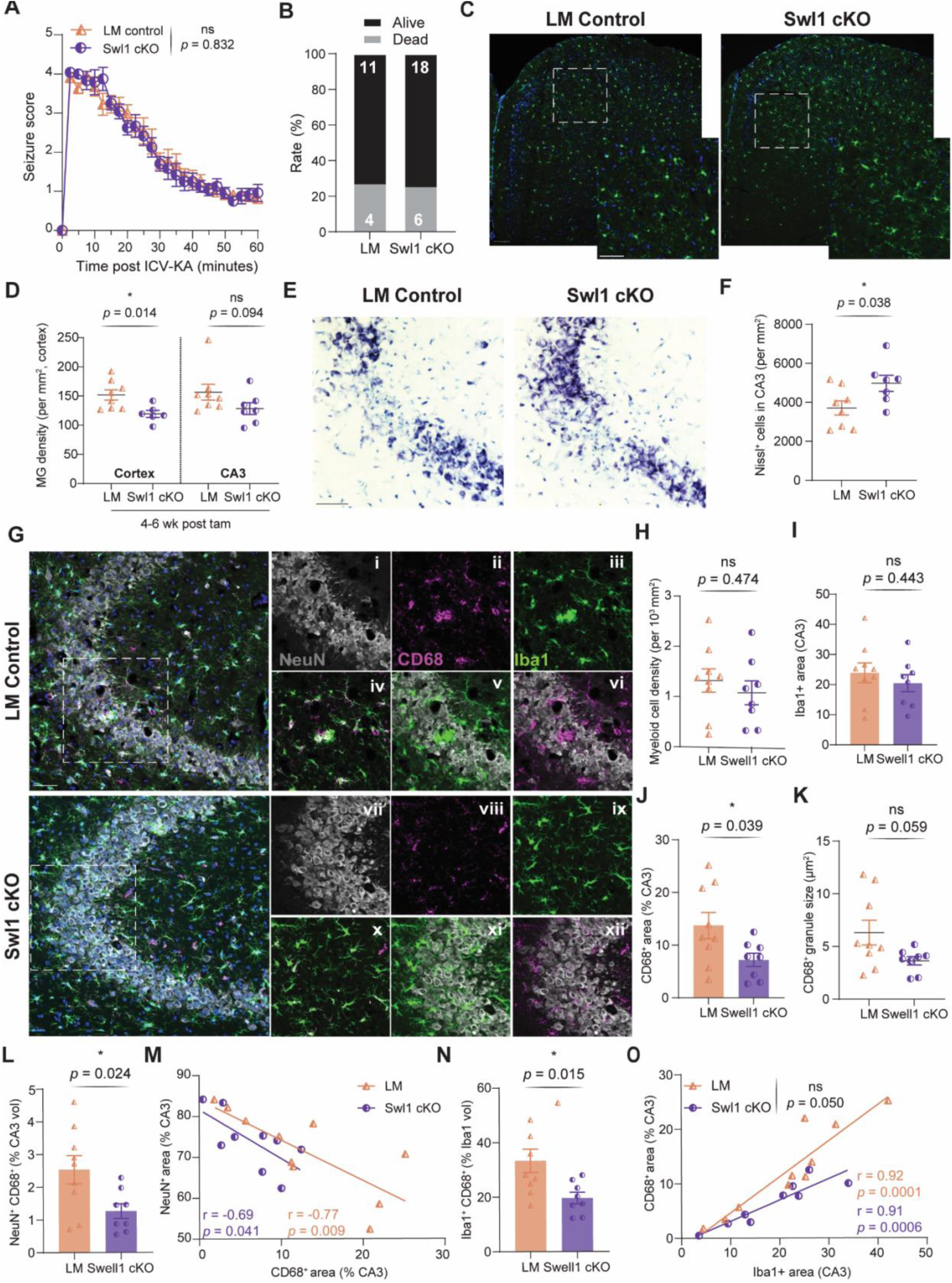
Female SWELL1 cKO mice exhibit neuroprotection without increased seizure severity. A) Longitudinal Racine seizure scores in Swell1 cKO vs littermate (LM) controls. (n=15-24 mice/group; statistic: 2-way ANOVA) B) Survival rate. C) Microglia density in cortex at 4-6 weeks after the last tamoxifen/ vehicle administration. Scale bar, 50 µm. D) Quantification of microglia density in cortex and CA3 (statistic: unpaired t-test and Mann-Whitney test respectively) E) Nissl staining showing neuronal loss in CA3 pyramidal layer at day 3 after seizure. Scale bar, 50 µm. F) Quantification of Nissl positive cell bodies at day 3 post seizure (statistic: unpaired t-test). G) Immunofluorescence images showing neuronal loss, myeloid and CD68 responses in CA3 at day 3 after seizure. Scale bar, 50 µm for images with overlay of all channels. H) (H-I) Quantification of myeloid cell density and Iba1^+^ area in CA3 pyramidal layer at day 3 after seizure (statistic: unpaired t-tests). J) (J-K) Quantification of CD68^+^ area and mean granule size in CA3 at day 3 after seizures (statistic: unpaired t-tests). L) Quantification of NeuN-CD68 double positive voxels in CA3 pyramidal layer (statistic: unpaired t-test). M) Correlation between CD68 ^+^ and NeuN ^+^ area in CA3 pyramidal layer at day 3 after seizure (statistic: Pearson correlation coefficient) N) Quantification of Iba1-CD68 double positive voxels in CA3 pyramidal layer as a percent of Iba1+ volume (statistic: unpaired t-test). O) Correlation between Iba1^+^ and CD68 ^+^ area in CA3 pyramidal layer at day 3 after seizure (statistic: Pearson correlation coefficient; simple linear regression to compare the slopes)

At 4–6 weeks post-tamoxifen, SWELL1 cKO females displayed a significant reduction in microglial density in the cortex and a non-significant decrease in the CA3 pyramidal layer (Fig. 6, C and D; Fig. S6C), mirroring the pattern seen in males. Furthermore, like males, female SWELL1 cKO mice exhibited reduced neuronal loss in CA3 three days after KA-induced seizures (Fig. 6, E and F). However, unlike males, female SWELL1 cKO mice did not show significant differences in the overall microglial response, as measured by Iba1⁺ area and cell density (Fig. 6, Giii, Gix, H, and I). Despite this, CD68 expression remained reduced in the CA3 region, with significantly lower CD68⁺ area and a non-significant reduction in CD68⁺ granule size (Fig. 6, Gii, Gviii, J, and K). These results indicate that lysosomal activation, rather than general microglial recruitment, may underlie the neuroprotective phenotype in females.

Consistent with this interpretation, SWELL1 cKO females showed reduced lysosome–neuron interaction, as evidenced by a lower volume of NeuN⁺ CD68⁺ voxels and a moderate (vs strong) negative correlation between NeuN⁺ and CD68⁺ areas (Fig. 6, Gvi, Gxii, L, and M).

Additionally, lysosomal synthesis efficiency appeared impaired, as shown by a reduced proportion of Iba1⁺ CD68⁺ voxels normalized to total Iba1 volume and a significantly flatter slope in the Iba1–CD68 correlation (Fig. 6, Giv, Gx, N, and O). These findings suggest that, as in males, impaired phagolysosomal processing may contribute to neuroprotection in female SWELL1 cKO mice.

To further explore sex-specific effects, we conducted a post hoc analysis comparing neuropathology responses across sexes and genotypes. No significant sex differences were observed in Iba1⁺ or CD68⁺ area in CA3 for either genotype (Fig. S6, D–F). However, females exhibited less neuronal loss than males, with statistically significant protection in SWELL1 cKO mice and a non-significant trend in littermate controls. Additionally, female littermate controls had a higher proportion of CD68⁺ Iba1⁺ voxels (normalized to Iba1⁺ volume) compared to males, despite similar slopes in the CD68–Iba1 correlation (Fig. S6, G–L). This difference was attributed to a higher intercept, suggesting that at any given level of microglial activation, females produce more lysosomal content. Biologically, this implies that while the rate of CD68 upregulation per unit Iba1 is similar across sexes, female microglia may exhibit a higher baseline lysosomal load.

## DISCUSSION

In this study, we investigated the role of microglial VRAC in acute seizures and post-seizure neuropathology, motivated by our observation of seizure-induced alterations in VRAC subunit expression. Using a conditional knockout model targeting the obligate VRAC subunit SWELL1 in myeloid cells, we uncovered three major findings: 1) SWELL1 deletion increased seizure severity in male mice, 2) it conferred paradoxical neuroprotection through impaired microglial phagocytosis, and 3) the effects varied by sex, with females displaying preserved neuroprotection despite unchanged seizure susceptibility. These results provide new insight into the role of microglial VRAC in modulating neuron–glia interactions in seizures and highlight a mechanistic link between ion channel function, neuroimmune activity, and sex-specific outcomes in epilepsy.

### Methodological considerations in microglial transgenic models for studies of epilepsy

Conditional LRRC8A/SWELL1 knockout models are essential for studying VRAC function, as global knockouts result in embryonic lethality, early postnatal death, and widespread organ dysfunction (*37*), underscoring the channel’s fundamental physiological importance. The discovery of the *Lrrc8* gene family and the identification of LRRC8A/SWELL1 as the obligate VRAC subunit (*24, 25*) have enabled the development of genetic tools to study its cell type– and context-specific roles. To date, four studies have directly examined microglia-specific VRAC function—three using conditional knockouts (*31, 38, 39*) and one employing AAV-mediated overexpression (*27*).

Some of these reports yield conclusions that diverge from both our findings and established VRAC literature in myeloid cells, necessitating careful methodological consideration. Efficient and specific microglial targeting remains challenging, even with recent advances in AAV vector engineering (*40*). For example, the AAV-mediated overexpression study did not assess transduction efficiency or specificity (*27*), limiting the interpretability of its results.

The conditional knockout studies have generally used CX3CR1-driven Cre expression to achieve microglia-specific deletion. Although CX3CR1 is the most widely used microglia-targeted Cre driver, it is also active in brain border-associated macrophages (BAMs) (*41–43*). Additional considerations with this system include: (1) the size of the loxP-flanked genomic region, which affects the likelihood of spontaneous recombination; (2) potential off-target effects of peripheral *CreER* activation; (3) tamoxifen toxicity; and (4) differences between constitutive and inducible Cre models. When studying microglia in seizure models, two other variables are critical: the genetic background of experimental groups, and CX3CR1 haploinsufficiency in Cre knock-in/knock-out lines. Genetic background can significantly affect seizure susceptibility and severity, even among closely related inbred strains (*44, 45*). Additionally, *CX3CR1* modulates both seizure severity and seizure-induced neuronal loss (*10, 46*), and its haploinsufficiency can impact microglial function (*10, 47, 48*).

In our model, the loxP-flanked region of the Swell1^(fl/fl)^ allele spans ∼5 kb (*49*), which minimizes the risk of spontaneous recombination (*41, 42*). We employed multiple control groups to account for experimental confounders: tamoxifen-injected non-littermate controls for CreER and tamoxifen-specific effects, and vehicle-treated littermate controls to account for CX3CR1 haploinsufficiency, genetic background, and residual recombination. Tamoxifen was administered at 5 weeks of age—a developmental time point that does not perturb the microglial transcriptome or morphology (*41, 50*)—followed by a 3–4 week recovery period to allow turnover of short-lived peripheral myeloid cells, thereby limiting recombination to long-lived tissue resident macrophages and microglia (*51*). We also excluded meninges, choroid plexus, and vascularized regions from our histological analyses to minimize BAM contamination, although some off-target recombination cannot be entirely ruled out. Future work may benefit from more microglia-specific genetic tools.

### How SWELL1 may influence microglia–neuron communication during acute seizures

Transcriptomic and proteomic analyses suggest that, at baseline, microglial VRAC may primarily be composed of LRRC8A (SWELL1) and LRRC8D subunits (Fig. 1B; Fig. S1C), consistent with previously published datasets (*11, 32, 52, 53*). Functional VRAC channels require LRRC8A in combination with at least one additional LRRC8 family member, with channel properties and substrate selectivity shaped by subunit composition (*23–25, 33*). For example, LRRC8D is necessary for the transport of neutral amino acids such as taurine and GABA, whereas LRRC8B and LRRC8C are dispensable for this function (*33*).

Following seizures, we observed downregulation of Lrrc8a and upregulation of Lrrc8b–d in microglia (Fig. 1, A and B), suggesting a potential shift in VRAC stoichiometry. This may reflect an adaptive response to the evolving demands of the post-seizure environment. However, since transcriptomic data were derived from CD11b⁺ populations, these changes could also reflect compositional shifts in myeloid subpopulations. To directly test the functional role of microglial VRAC in seizures, we generated a SWELL1 cKO model in microglia, using an experimental design optimized to minimize peripheral effects and isolate microglia-specific contributions.

SWELL1 cKO males exhibited increased seizure severity (Fig. 1, F–I). To investigate underlying mechanisms, we focused on two VRAC-associated processes: regulation of cell proliferation and release of neuroactive metabolites (*33, 37*). We observed a transient reduction in microglial density in the cortex and CA3 region during the time window in which seizures were induced (Fig. 2, A–D; Fig. S1, D–G). This contrasts with two previous studies that found no baseline differences in microglial density in SWELL1 cKO models (*31, 38*). Potential explanations for this discrepancy include the use of constitutive versus inducible Cre drivers, limited sample size in prior studies, and the transient nature of the effect, which resolved by 10 weeks post-tamoxifen treatment in our model.

Microglial depletion has previously been shown to exacerbate seizure severity in the kainate model (*6, 7*), raising the possibility that even a modest (∼10–20%) reduction in microglial density could contribute to heightened excitability. However, we found no differences in microglia–neuron physical interactions two hours after seizure (Fig. 2, H and I; Fig. S2, G and H). This may be due to compensatory morphological changes such as increased microglial territory size (Fig. 2F), which could allow fewer microglia to maintain normal neuronal surveillance.

Interestingly, despite an overall reduction in soma size from baseline after acute seizures, SWELL1 cKO microglia were larger than those in controls (Fig. 2, J–L). Prior studies have established that VRAC activation triggers the release of various organic and inorganic anions, including inhibitory and excitatory neuromodulators such as taurine, GABA, and glutamate (*24–26, 33, 54*). We hypothesized that impaired release of inhibitory metabolites might disrupt the excitatory/inhibitory balance and contribute to increased seizure severity (Fig. S3C). Release of taurine, GABA, and glutamate was significantly attenuated in SWELL1 cKO microglia (Fig. 3, B–D) while ATP was unaffected (Fig. 3G), contrasting with reports from BV2 and HeLa cells (*31, 55*). This discrepancy may reflect differences in cell type, VRAC composition, or activation conditions (e.g., sphingosine-1-phosphate vs. hypotonic stress) (*31–33, 55*). We extended these findings in vivo by analyzing CSF collected from mice before and one hour after seizure induction (Fig. 3H). While baseline levels were similar across groups, SWELL1 cKO mice exhibited lower post-seizure levels of taurine and glutamate, but not GABA (Fig. 3, I–K; Tables S2–S3). Further, dysregulated taurine homeostasis was seen in SWELL1 cKO animals, (Fig. S3, D and E). These changes may reflect loss of microglial SWELL1, differences in seizure severity, or an interaction between the two.

It is important to note that CSF metabolite concentrations may not accurately reflect those in brain interstitial fluid (*56*), particularly during seizures when microglial processes physically interact with neurons (*7, 9, 57*). Such proximity could locally concentrate VRAC-released neuromodulators. Taurine, for example, is a weak GABA-A receptor agonist with established anticonvulsant effects (*58–61*), and its reduction could reduce inhibitory tone. However, glutamate was also reduced in SWELL1 cKO CSF post-seizure. A previous study found elevated seizure susceptibility and mortality in mice with brain-wide SWELL1 deletion (under a Nestin-Cre driver) despite reduced tissue glutamate levels (*62*), suggesting that VRAC disruption may produce complex shifts in excitatory/inhibitory balance. These findings warrant further investigation using real-time, spatially resolved metabolite sensing in intact brain tissue.

### Phagocytic defects in SWELL1-deficient microglia underlie neuroprotection following seizures

Despite experiencing more severe seizures, SWELL1 cKO mice exhibited reduced neuronal loss in the CA3 region of the hippocampus (Fig. 4, A to D). CA3 is particularly vulnerable to kainate-induced seizures due to its enrichment in kainate-type glutamate receptors and central role in limbic seizure circuitry (*11, 12, 16, 63, 64*). Previous studies have identified phagoptosis—a form of cell death driven by microglial engulfment of stressed-but-viable neurons—as a key contributor to post-seizure neurodegeneration (*13, 21, 65*). This prompted us to investigate whether altered phagocytic responses in SWELL1 cKO mice could account for the observed neuroprotection.

Following seizures, we observed reduced microglial proliferation in SWELL1 cKO males(Fig. 4, B, E, and F), consistent with prior findings in models of neuropathic pain (*31*). Lysosomal responses, as assessed by CD68 immunostaining, were also diminished in SWELL1 cKO animals (Fig. 4, B, G–I). CD68 is a member of the LAMP protein family and a robust marker of lysosomal expansion following seizure-induced neuronal injury (*11–13, 16, 66*). NeuN⁺ CD68⁺ voxels—interpreted as neuronal material within or interacting with phagolysosomes were significantly reduced in SWELL1 cKO mice, (Fig. 4, J and K), supporting impaired phagocytosis as a driver of neuroprotection. Further, the reduction in lysosomal response in SWELL1 cKO animals was independent of the low myeloid response as indicated by a lower volume of Iba1⁺ CD68⁺ voxels, normalized to total Iba1⁺ volume (Fig. 4, N and O). Although CD68 and Iba1 areas were strongly correlated in both groups, the slope of this relationship was significantly reduced in SWELL1 cKO mice (Fig. 4P; Fig. S4J), suggesting reduced lysosomal generation per unit of myeloid mass. Together, these data support impaired lysosomal biogenesis as an intrinsic deficit in SWELL1-deficient microglia.

The mechanisms by which VRAC regulates lysosomal biogenesis remain unclear. One possibility is a direct role for VRAC in lysosomal volume regulation. A recent study demonstrated that lysosomal VRAC currents modulate lysosome size and homeostasis in response to cellular stress (*67*), supporting a potential direct role. Alternatively, the defect may arise from impaired upstream processes such as phagocytic engulfment, which influence downstream lysosomal load. *In vitro* studies revealed that proliferative and phagocytic defects observed in SWELL1 cKO microglia were cell-autonomous and that engulfment kinetics was reduced in addition to phagolysosomal defects. Collectively, our results indicate that SWELL1 regulates both microglial proliferation and phagocytosis in a cell-intrinsic manner. In the context of seizures, these impairments blunt phagocytosis and thus mitigate secondary neuronal loss— providing a mechanistic explanation for the paradoxical neuroprotection observed in SWELL1 cKO mice despite worsened seizures in male mice.

### Sex specific effects of microglial VRAC

Sexually dimorphic roles of microglia have been described in neuropathic pain and SWELL1 cKO models (*31, 34*), supporting the plausibility of such interactions in epilepsy. Notably, female SWELL1 cKO mice did not exhibit increased seizure severity (Fig. 6, A and B; Fig. S6, A and B). Previous studies have demonstrated that microglial depletion exacerbates both acute and chronic seizures in male mice (*6, 7*), but similar investigations have not been conducted exclusively in females. Although sex differences in kainate-induced seizure models are well established (*68*), the role of microglia in mediating these differences remains largely unexplored. As such, it is unclear whether the sex-dependent effects observed in our model reflect a broader microglial dimorphism or a VRAC-specific phenomenon. One possibility is that taurine release from microglia during seizure may be a male specific response, dependent on VRAC subunit composition and will be examined in future studies.

With respect to post-seizure neuropathology, female SWELL1 cKO mice showed no significant differences in Iba1⁺ microglial responses relative to littermate controls, in contrast to the attenuated microgliosis observed in males (Fig. 6, G–I). Prior work suggests that female mice exhibit less robust microgliosis than males following kainate-induced seizures (*68*), and it is possible that our 3-day post-seizure time point was too early to detect such changes in females. Nonetheless, our findings raise the possibility that SWELL1 influences microglial proliferation or activation in a sex-dependent manner. Importantly, like males, female SWELL1 cKO mice exhibited reduced CD68 expression, diminished lysosomal biogenesis, and attenuated neuronal loss (Fig. 6, E–G, J–O), underscoring a conserved role for VRAC in regulating microglial phagocytic capacity. These findings suggest that SWELL1-dependent modulation of microglial phagocytosis may be a shared mechanism of neuroprotection across sexes, whereas its role in seizure susceptibility may be male-biased. Such differences may inform precision medicine approaches: for instance, microglia-targeted anti-SWELL1 therapies might offer neuroprotective benefits in female epilepsy patients without increasing seizure risk.

### Future directions

Our future investigation will focus on identifying the mechanisms responsible for VRAC activation in the ictal and post-ictal phases. Some of the candidates include osmolarity changes, increased ATP, intracellular calcium, and superoxide production, all of which known VRAC activators and observed in ictal phase (*72–75*). Similarly, during the days and weeks following the seizure, sphingosine-1-phosphate (S1P) may drive VRAC activation. S1P is a known VRAC activator (*23, 72*) as well as a ‘find me’ phagocytic signal, acting via the S1P receptor family (*21*). It is possible that some of the phagocytic activity attributable to S1P may be driven by VRAC rather than S1P receptors and needs to be examined.

## Author contributions

Abhijeet S. Barath*: conceptualization, study design, conducted experiments, data analysis, writing manuscript, drafting figures

Aastha Dheer*: conceptualization, study design, conducted experiments Emily Dale: conducted experiments

Flavia Goche: conducted experiments

Thanh Thanh Le Nguyen: conducted experiments, data analysis, drafting figures Laura Montier: conducted experiments, data analysis

Mastura Akter: conducted experiments Mekenzie Peshoff: conducted experiments FangFang Qi: experimental design Anthony Umpierre: study design

Dale B. Bosco: experimental design, generated SWELL1^(fl/fl)^: CX3CR1^(CreER/CreER)^ and SWELL1(fl/fl): tdT(fl/fl) mice

Koichiro Haruwaka: provided initial image J analysis scripts, experimental designs Rajan Sah: provided SWELL1^(fl/fl)^ mice, conceptualization

Long-Jun Wu: conceptualization, study design, writing manuscript, funding

## Acknowledgements

We acknowledge the invaluable contributions of following individuals and cores for their support: 1) Aswath Chandrasekhar (Department of Pathology, Mayo Clinic), August John (Department of Molecular Pharmacology and Experimental Therapeutics, Mayo Clinic), as well as all members of the Wu lab for their insightful discussions; 2) Mayo Clinic metabolomics core (Mai T Petterson, Jaime Gransee, and Peggy S. Helwig), Mayo Clinic microscopy core (Duane M. Deal and Alex A. Anwar), UTH-Houston IMM Microscopy core (Zhengmei Mao and Ryan Durham) for providing technical expertise and services, and Caiyun Liu at Mayo Clinic for her assistance with some of the experiments.

## METHODS

### Animals

SWELL1^(fl/fl)^ mice were kindly provided by Dr. Rajan Sah of Washington University at St. Louis. CX3CR1^(CreER/CreER)^ mice (B6.129P2(Cg)-*Cx3cr1^tm2.1(cre/ERT2)Litt^*/WganJ; strain # 021160), tdT^(fl/fl)^ mice (B6.Cg-*Gt(ROSA)26Sor^tm14(CAG-tdTomato)Hze^*/J; Strain # 007914), and wid type mice (C57BL/6J; Strain # 000664) were purchased from the Jackson lab. SWELL1^(fl/fl)^:tdT^(f/f)^, SWELL1^(fl/fl)^:CX3CR1^(CreER/CreER)^, and SWELL1^(fl/fl)^:tdT^(fl/-)^:CX3CR1^(CreER/wt)^ (Table S6) mice were generated in house through crossbreeding. Both sexes were used. They were housed in temperature and humidity-controlled environments with 12-hour day-night cycles and ad-libitum access to food and water. To induce gene recombination, tamoxifen (150 mg/kg) or corn oil (vehicle; Table S6) was administered intraperitoneally to 5-week-old mice on alternate days for a total of 4 injections. This resulted in three groups of experimental animals-Seizure experiments were performed at least 3-4 weeks after the last tamoxifen/corn oil injection. All experimental procedures were approved by the Institutional Animal Care and Use Committee (IACUC) at Mayo Clinic, Rochester, MN (protocols A00002290-16-R22 and A00002731-17-R23) and the Center for Laboratory Animal Medicine and Care (CLAMC) at UTHealth Houston (protocols AWC-24-0004 and AWC-24-0046).

| Designation | Genotype | Substance administered | Rationale |
| --- | --- | --- | --- |
| SWELL1 cKO | SWELL1<sup>(fl/fl)</sup>: tdT<sup>(fl/-)</sup>: CX3CR1<sup>(CreER/wt)</sup> | Tamoxifen | Induce myeloid specific conditional deletion of SWELL1 |
| Littermate (LM) controls | SWELL1<sup>(fl/fl)</sup>: tdT<sup>(fl/-)</sup>: CX3CR1<sup>(CreER/wt)</sup> | Vehicle (corn oil) | Control for genetic background and CX3CR1 heterozygosity |
| Tamoxifen (Tam) controls | SWELL1<sup>(fl/fl)</sup>: tdT<sup>(fl/fl)</sup> or tdT<sup>(fl/-)</sup>: CX3CR1<sup>(CreER/wt)</sup> | Tamoxifen | Control for adverse/non-specific effects of tamoxifen |

### RNA sequencing data and analysis

Microglial bulk RNA sequencing data recently published by us (*11*) and publically available was reanalyzed to assess changes in microglia homeostatic genes, known seizure related genes, and Lrrc8 family in WT mice before and 7 days after seizure. Expression changes are presented as Log2 fold change or FPKM (Fragments Per Kilobase of exon per Million mapped fragments) values and significance was assessed with *q values* (false discovery rate adjusted *p values*).

### Microglia isolation from adult brain and qPCR

Microglia isolation protocol was adapted from Scheyltjens., et al., 2022 (*76*). Briefly, mice were deeply anesthetized, perfused with ice-cold PBS, and brains were rapidly removed. Olfactory bulbs and cerebellum were discarded. The remaining cerebrum was cut into fine pieces, transferred to 6-well plates containing 2 ml RPMI media/well, and maintained on wet ice.

Enzymatic (cocktail of DNAse I, collagenase I, and collagenase IV) and mechanical (pipette trituration) dissociation of the tissue were performed followed by straining (70 µm filters), and removal of myelin and debris using a 30% Percoll gradient. Cells were then pelleted with centrifugation, resuspended, and magnetically labelled with the EasySep™ Mouse CD11b Positive Selection Kit II. This kit targets CD11b+ cells for positive selection with antibodies recognizing the CD11b surface marker. Following manufacturer’s instructions, both CD11b positive (predominantly microglia) and negative (astrocytes, oligodendrocytes, few neurons) fractions were collected. CD11b positive fraction was used to evaluate SWELL1 cKO efficiency, while CD11b negative fraction was used to assess the specificity.

Cells were then lysed using RNeasy® Micro Kit and RNA was extracted following manufacturer’s instructions. RNA quantity was measured with microvolume spectrophotometer. It was then reverse transcribed to cDNA using iScript™ cDNA Synthesis Kit following manufacturer’s protocol. Real time quantitative PCR for Lrrc8a (Swell1) and GAPDH were then performed using SsoAdvanced Universal SYBR Green Supermix. Details of all the reagents may be found in Table S6.

### Induction of kainate status epilepticus

9-12-week-old mice were provided with ibuprofen (0.2 mg/ml) analgesia in their drinking water 3 days prior to their surgery. They were induced with 3% isoflurane and maintained on 1.5-2% with oxygen flow during surgery. Skull was exposed and a 0.9-1 mm wide hole was stereotactically drilled, centered at 1 mm lateral and 0.2 mm posterior to bregma on the left side. A 6 mm long, 25 G stainless steel cannula (Table S6) was then implanted in the left lateral ventricle with its bottom at 2.05 mm below the skull surface. The cannula was sealed in placed and the incision was covered with layered application of iBond and dental cement (Table S6) and cured with UV light for 1 and 2 minutes respectively. Ibuprofen water was discontinued 3 days after surgery. On day 4, 0.1-0.12 µg kainate (Table S6) dissolved in 4-5 µl of PBS was injected through the cannula to induce acute seizure. The mice were then placed in clear cylindrical buckets and observed for 2-3 hours. All seizure experiments, except a few early cohorts were recorded using a GoPro camera.

A seizure score based on the modified Racine scale (*9*) was assigned every 5 minutes. Briefly the following scoring matrix was used-(1) freezing behavior, (2) frozen with tail raised, (2.5) tail raised and lying on its side, unable to maintain posture, (3) reared up with forepaw and/or head clonus, (3.5) forepaw and/or head clonus while lying on the side, unable to maintain posture, (4) rearing and falling, severe clonic movement of multiple extremities, or stereotypic running movements with periods of stillness, (5) level 4 activity exceeding 15 minutes, vigorous jumping, (6) tonic phase arrest (extensor tonic posturing of both upper and lower limbs and ventral flexion of tail) with brief cessation of respiration followed by spontaneous return to fast breathing, (7) tonic phase arrest with irreversible cessation of respiration and death. Only those mice who achieved a seizure score >3 within the first 30 minutes of ICV-KA administration were included for further analysis. All seizure phenotyping was performed in a genotype blinded manner to prevent observer bias. Longitudinal and average seizure scores, proportion of mice experiencing at least one severe seizure (score ≥ 5), and mortality were compared between genotypes.

### Tissue collection for histopathology

Mice were deeply anesthetized with isoflurane and sequentially perfused transcardially with 30 ml each of ice-cold PBS and 4% paraformeldehyde (PFA. Brains were rapidly removed and post-fixed in 4% PFA for 24 hours followed by 30% sucrose in PBS for at least 4 days for cryoprotection. After freezing in molds with OCT, 20 um thick coronal sections from anterior to posterior hippocampus were cut using a cryostat and placed on adhesive glass slides. The slides were stored at −20 °C until further use. Details of all the reagents may be found in Table S6.

### Immunofluorescence staining

Slides were warmed to room temperature and washed with tris buffered saline (TBS). Permeabilization and blocking was performed with 10% donkey or goat serum prepared in 0.4% Triton X-100 in TBS for one hour (Table S6). Slides were then incubated overnight (at least 12 hours) at 4 °C with primary antibodies prepared in 5% donkey or goat serum in 0.4% Triton X-100 solution in TBS. Please see Table S6 for a list of primary antibodies and concentrations used. After overnight incubation, the slides were allowed to warm up to room temperature for an hour followed by 3 washes with 0.1% Triton X-100 prepared in TBS. They were then incubated at room temperature for 1.5-2 hours with corresponding fluorophore conjugated secondary antibodies prepared in 0.4% Triton X-100 solution in TBS. After a few rounds of washing with 0.1% Triton X-100 prepared in TBS slides were counterstained with mounting DAPI and coverslipped. Slides were allowed to dry 24 hours before imaging or storage (−20 °C) for future imaging.

### Microglia density analysis

To determine the density of microglia, 3-5 sections per region were imaged with Nikon TE2000e fluorescence microscope for each animal. Single z-plane, multi-channel images for were taken with a 10x objective for cortex (FOV: 1331.2 x 1328.6 µm^2^) and 20x objective for the CA3 region (FOV: 665.6 x 664.3 µm^2^). ImageJ was used to analyze the images using a code with the following logic sequence-(1) Preprocess DAPI: rolling background subtraction with radius 10-20 pixels (larger than the largest foreground object of interest) to suppress autofluorescence; auto threshold with Otsu method. (2) Ask user to draw the boundaries of the region of interest (cortex or CA3 ROI) on a copy of DAPI channel. (3) Preprocess Iba1 channel: 2-step background subtraction-first, suppress any areas with holes (missing tissue) by creating a mask of pixels below the autofluorescence threshold (empirically determined as <3 mean gray value for 8-bit images) and subtract the masked area from main image; second, rolling background subtraction with radius 20-30 pixels to suppress autofluorescence; auto threshold with Otsu method. (4) Image calculator was used to find DAPI^+^ Iba1^+^ microglia nuclei on thresholded images using ‘AND’ function. (5) Area and centroid functions were enabled in the set measurements menu. (6) On the result of image calculator apply the cortex or CA3 ROI and use ‘analyze particles’ function to obtain a count of Iba1^+^ DAPI^+^ particles above 15 um^2^ in size. (8) Microsoft excel was used to compute density based on the particle count in the ROI and the ROI area. (9) NND plugin was used to obtain the nearest neighbor distances based on the centroid values for microglia nuclei. tdTomato channel was analyzed in the same manner as Iba1.

### Iba1^+^, CD68^+^, and NeuN^+^ area analysis

Sections were imaged and analyzed in the same way as for microglia density analysis with the following changes in the logic sequence-(1) Triangle (instead of Otsu) method was used to threshold the Iba1 channel as it was found to be more sensitive to finer processes. (2) All Iba1 positive areas above the threshold value within the ROI were summed and included for analysis without a size filter. (3) Microsoft excel used to compute % of ROI area positive for Iba1.

### Microglia territory and Sholl analysis

Iba1 stained sections were images with a Zeiss LSM 780 confocal microscope at 20x magnification (FOV: 424.26 x 424.26 µm^2^), visually centered on the CA3 region. A z stack volume was then acquired at 1024×1024 pixel resolution with a 1 μm step size across a uniform 10 μm stack. ImageJ was used to analyze the images using a code with the following logic sequence-(1) Iba1 volume was z-projected with ‘max-intensity’ option and two copies were made. (2) First copy was auto-thresholded with triangle method. (3) Contrast was enhanced in the second copy with saturation of 1% of the pixels for improved visualization of finer processes. (4) DAPI volume was z-projected with ‘max-intensity’ option and CA3 ROI was drawn and transferred to the contrast enhanced copy of the Iba1 channel. (5) Microglia with somas within the CA3 were identified and boundaries were drawn for their territory (MG territory ROIs). (7) MG territory ROIs were transferred to the thresholded images, and ‘multi-measure’ operation of the ROI manager was used to determine the territory sizes. (8) An ImageJ plugin(*77*) was used to calculate the Sholl morphology of individual microglia. The center of the soma was user defined, and intersections were calculated along the path of concentric circles starting 5 μm from the soma center and increasing by 1 μm in radius.

### Volumetric interaction analysis

Tissue sections stained with multiple markers were imaged with Nikon AXR confocal microscope with either-(a) 40x water immersion objective (FOV: 435.35 x 435.35 µm^2^) at 2048×2048 pixel resolution with 0.5 µm step size across a uniform 10 µm stack, or (b) 20x objective (FOV: 878.5 x 878.5 µm^2^) at 2048×2048 pixel resolution with 1 µm step size across a uniform 10 µm stack. Setting used for a given experiment may be found in the results section. Volumetric interaction analysis was performed with imageJ using the following logic sequence: (1) Image volume was split into channels. (2) Stack thresholding was performed for each channel with ‘stack histogram’ enabled-(a) Otsu method for DAPI; (b) Li method for NeuN; (c) Triangle method for Iba1; (d) Triangle method for P2Y12; (e) Fixed threshold with bounds 25-255 (8-bit images) for CD68. (3) CA3 ROI was drawn on z-projected DAPI volume (max-intensity method). (4) ‘AND’ function of image calculator was applied to a pair of thresholded channel volumes of interest to obtain a result stack that’s double positive for both channels. (5) CA3 ROI was transferred to the result stack. (6) ‘Analyze particles’ function with no-size filter was applied to obtain a layer-by-layer area of the result stack within the CA3 ROI. (7) Area was multiplied by voxel thickness (0.5 µm for 40x images, 1 µm for 20x images) and summed to obtain the imaging volume positive for the chosen pair of channels of interest. (8) Double positive channel volume were then divided by the total CA3 volume (CA3 ROI area x total stack thickness of 10 µm) or Iba1 channel volume for normalization. Note: fixed bounds of threshold were used for CD68 due to poor performance of auto thresholding methods in sections with very low CD68. The bounds were empirically determined by visual inspection of parameter performance on the brightest and least bright sections of CD68 channel (max intensity z-projected images) among all the sections of all male animals examined.

### Microglia soma size analysis

Soma size analysis was performed on Iba1 stained sections imaged with a Zeiss LSM 780 confocal microscope at 20x magnification (FOV: 424.26 x 424.26 µm^2^), 1024×1024 pixel resolution with a 1 μm step size across a uniform 10 μm stack, centered on CA3. Using ImageJ, Iba1^+^ DAPI^+^ microglia nuclei within CA3 were found using the same logic sequence as described in microglia density analysis. Following additional steps were performed to obtain true soma-(1) Max-intensity z projected Iba1 images were auto thresholded using Otsu method. (2) Two iterations of ‘open’ function (process menu → binary → options) were applied to get rid of most of the microglia branches. (3) Four iterations of ‘dilate’ function (process → binary → dilate) were applied to the Iba1^+^ DAPI^+^ thresholded microglia nuclei to add material around them. (4) Image calculator with ‘AND’ function was applied to the results of steps (2) and (3) to obtain true soma, devoid of almost all branches. (5) Area and shape descriptors were enabled in set measurements menu. (6) Analyze particles operation was performed with a size filter of 17 µm^2^. (7) Circularity and area were noted to obtain the soma size and activation status respectively. ImageJ calculates circularity using the formula, circularity = 4 π (area/perimeter^2^) where 1 indicates a perfect circle and values approaching 0 indicate increasingly elongated polygon.

### Primary microglia cultures

Primary microglial cultures were prepared from brains of neonatal mice (Fig.S3A). Briefly, brains were extracted from 2–7-day old pups. Cerebellum, olfactory bulb, and meninges were removed. The brains were then mechanically and enzymatically digested into single cell suspension using 0.25% trypsin. The suspension was filtered through a 100 µm sterile strainer. The resulting mixed glial suspension was cultured in DMEM:F12 media supplemented with 10% fetal bovine serum and 1% penicillin/streptomycin in T-75 flasks for 12-17 days. Once the astrocytic layer reached confluence and abundant microglia were visualized in the mixed cultures (round, pearly cells loosely adherent to the top of astrocytes or floating in the media), they were treated with 5 uM 4-hydroxy tamoxifen (4-OHT) or vehicle (DMSO) for 48 hours. To separate microglia, flasks were shaken at 200 rpm for 30 minutes. Microglia density was counted using a hemocytometer and trypan blue stain, and they were plated in d-lysine coated 12-well or 24-well culture plates (at density ranging from 30,000-50,000 cells per well). Experiments were performed 48 hours after replating purified microglia. Microglia were maintained on astrocyte conditioned DMEM:F12 media. Details of all the reagents may be found in Table S6.

### *In vitro* metabolomics assay

Primary microglia cultures were washed thrice with isoosmotic PBS to remove all traces of the media. They were incubated in isotonic aCSF (140 mM NaCl, 5 mM KCl, 1 mM MgCl2, 1 mM CaCl2, 10 mM HEPES, and 5 mM glucose; osmolarity was adjusted to be between 300-320 mOsm/L, pH adjusted to 7.3-7.4; 350 µl/well of 12 well plate) at 37 °C (*31*). The isotonic buffer was carefully removed to a pre-labelled 1.5 ml conical vial and cells were incubated in a 30% hypotonic aCSF (7 parts of isotonic buffer diluted with 3 parts of distilled water; osmolarity 200-215 mOsm/L, pH 7.3-7.4; 350 µl/well of 12 well plate) for 30 minutes at 37 °C to activate VRAC currents. Hypotonic buffer was also removed to a 1.5 ml vial. All vials with samples were spun down at 4000 g for 5 minutes to pellet any cells. Supernatants were carefully removed to another set of labelled vials inside a laminar flow cabinet and sealed shut. Samples along with two vials, each of unused isotonic and hypotonic solutions (negative controls) were stored at −80°C or immediately transported on dry ice to the Mayo Clinic Metabolomics Core for analysis. Liquid chromatography/ mass spectrometry (LC/MS) was used to analyze fourteen neuromodulator panel metabolites. Average values of respective isotonic and hypotonic negative controls were subtracted from sample values before statistical comparison.

### Colorimetric assay for ATP

Isotonic and 30% hypotonic stock solutions were prepared as discussed above with the addition of 100 µM ARL 67156 trisodium salt. ARL salt is an ectonucleotidase inhibitor and reduces hydrolysis of ATP. The metabolomics assay was repeated with the ARL added buffers.

Following manufacturer’s instructions, ATP standards were prepared separately for isotonic and hypotonic buffers in a range of 10 pM to 10 nM. ATP assay mix was loaded into a 96 well plate and rested for 3 minutes to hydrolyze endogenous ATP. Samples were added to the ATP assay mix and pipetted a few times. Luminance was read immediately on a spectrophotometric plate reader. Sample ATP values were derived from the standard calibration curves.

### CSF and plasma collection experiments

CSF and plasma were collected in different groups of animals at baseline (pre-seizure) and 1-hour after ICV-KA induced status epilepticus. Briefly, mice were induced with 3% isoflurane and maintained on 1.5% with oxygen flow. Their head was flexed forward to expose the back of the neck. A vertical incision was made from just behind the ears to upper thoracic level. Neck muscles were carefully dissected to expose the atlantooccipital membrane overlying the cisterna magna. Blood was wiped with sequential application of wet (PBS) and dry sterile cotton tips.

This step is vital to avoid any blood contamination. Under a surgical microscope, a sharp tipped pipette prepared from a glass capillary (Table S6) was guided towards the atlantooccipital membrane using a micromanipulator. The tip was pointed just lateral to the midline, away from visible blood vessels. The capillary was gently taped at the back to facilitate entry into the cisternal space. Its position was readjusted until clear CSF flow was obtained. Around 2-8 µL of CSF was collected per mouse. For post-seizure group, CSF was collected for only 5-6 minutes to minimize the impact of anesthesia on CSF metabolites. CSF was then diluted with PBS in a 1:24 or 1:9 ratio to obtain the minimal volume of 50 µL required for LC/MS analysis as per core specifications. Any samples with visible blood contamination or <1.5 µL volume were discarded. Samples were stored on wet ice until further processing. Following CSF collection, mice were deeply anesthetized by isoflurane and thoracotomy was performed. 200-300 µL blood was obtained directly from the left ventricle using a 25G needle and syringe and transferred to vials containing EDTA. This was followed by perfusion with ice-cold PBS, 4% PFA, and brain collection. Vials with blood and CSF were centrifuged at 8000 g for 10 minutes to pellet cells.

80-100 µL of yellowish to clear plasma supernatant was carefully pipetted to a 0.5 ml conical vial. Similarly, available volume of CSF supernatant was pipetted into new vials. CSF and plasma vials were stored at −80 °C until transport to the Mayo Clinic Metabolomics Core for analysis. PBS used for diluting CSF served as negative control. Average values of negative controls were subtracted from sample values followed by correction for dilution before statistical comparison. Metabolite values were normalized to controls in each batch to reduce batch to batch variability.

### Nissl (Cresyl violet) staining and analysis

Nissl (Cresyl violet or CV) staining was performed to assess density of healthy neurons at baseline at 3 days after seizure. CV stains Nissl substance in the cytoplasm of neurons in PFA or formalin-fixed tissue. Previously cryosectioned 20 µm thick coronal sections were allowed to warm up to the room temperature. Sections we defatted with xylene, hydrated with serially decreasing concentrations of ethanol solutions, washed with tap water, and then incubate with freshly prepared, acidified (pH 3.5-3.8) 0.1% CV solutions for 10 minutes at 37 °C. They were differentiated with a glacial acetic acid solution in 95% ethanol, and brief successions of increasing concentrations of ethanol. Finaly, sections were dehydrated with 100% ethanol, cleared with xylene and coverslipped using depex mounting media. After drying for 24 hours in a fume hood, imaging was performed with Zeiss Axio inverted microscope, centered on CA3.

For analysis using ImageJ, CA3 boundaries were drawn on the image and the number of healthy appearing neuronal somas were marked using the ‘multi point’ tool. Non-viable neurons identified as small-sized, dense, irregular-shaped pyknotic cells were excluded. Density of viable neurons within the CA3 ROI was computed. Details of all the reagents may be found in Table S6.

### *In vitro* microglia density analysis

Primary microglia separated from mixed culture were imaged at day 10 after replating with Keyence BZ-X800 fluorescence microscope at 20x magnification. Single z-planer images were obtained with TRITC (tdTomato) and phase contrast filters. ImageJ was used to quantify the number of tdT^+^ cells in single fields of view per well by auto thresholding with Shanbag method and watershed (process → binary → watershed).

### In vitro phagocytosis assay

Aqueous green fluorescent latex beads (Table S6) were opsonized by incubating in FBS in a 1:5 volume by volume ratio for 1 hour at 37 °C. This coats the bead with IgG which enhances phagocytic activity of microglia via the Fcγ receptors. The bead-FBS mixture was diluted with serum-free DMEM:F12 to reach a volume/volume concentration of 1:5:1994 for beads: FBS: media. It’s very important to use serum free DMEM:F12 for this dilution step as serum IgG would saturate the microglia Fc receptors, significantly reducing phagocytosis of opsonized beads. Primary microglia were washed with warm PBS 2-3 times followed by addition of serum free DMEM:F12 with opsonized beads. Imaging was performed at 5 minutes intervals on Keyence BZX800 microscope (with added incubator module), or after for 1-2 hours in a regular incubator at 20x magnification. Microglia were visualized using endogenously expressed red tdTomato signal (TRITC filter) while the beads were visualized using GFP filter. Phagocytosis was quantified as the percentage of microglia with phagocytic inclusions and the average phagocytic load per microglia (percentage of individual microglia area positive for GFP).

### Modified regulatory volume decrease assay

Purified primary microglia were washed 2-3 times with PBS and incubated in 700 µl isotonic aCSF per well of a 12-well plate. Single z-planer phase contrast images were obtained every two minutes with Keyence BZX800 microscope (with added incubator module) at 20x magnification. After 10 minutes of imaging in isotonic aCSF, 300 µL of distilled water was gently added along the margins of wells to create 30% hypotonic solution. Imaging was continued every two minutes for another 30 minutes. Soma size was manually quantified using imageJ and compared at various time points between the experimental groups.

### Assessment of culture purity

Culture purity was quantified as the percent of tdTomato positive cells among all the cells observed with phase contrast microscopy in the field of view (Fig.S3B).

### Statistics, rigor, and reproducibility

Sample sizes were determined using G*Power 3.1 (*78*) based on pilot experiments conducted on 5-10 animals per group, aiming to minimize animal usage while ensuring sufficient sample to detect robust effect sizes. False discovery rate (FDR) adjusted *p-values* (aka *q values*) were reported for RNA sequencing data. In general, data from 3-5 hippocampal or cortical sections were averaged per animal to obtain histopathology values for comparison. Normality was assessed (where applicable) using the Shapiro-Wilk test, chosen for its effectiveness for both small and large sample sizes. Normally distributed data were analyzed using unpaired t-tests while Mann-Whitney (nonparametric) test was used otherwise. Chi-square analysis was used to compare proportions. 2-way ANOVA with Bonferroni’s multiple comparison test was used for data with more than two timepoints from the same animal or culture sample to compare genotype x time/ condition effects. 2-way ANOVA was also used to compare the distribution of properties like Sholl intersections x radius and nearest neighbor distances x % microglia between genotypes. Nested t-tests were used when individual cell properties like microglia territory sizes were compared to account for variation that might be present within individual animals, making the comparison of the main genotype effects more robust. Correlations were examined using Pearson r. Simple linear regression was used to compare the correlation slopes. CSF metabolite data were normalized to the average of respective littermate controls per experimental batch to correct for batch-to-batch variability. Permutation test was used to compare the normalized CSF metabolite data for a more rigorous accounting of small sample sizes, non-standard distributions, and residual batch effects not corrected by normalization. Raw values for all metabolites for all experiments and groups are provided in the supplementary tables.

Experiments were designed based on well-established protocols published in literature and validated by the authors. All seizure experiments were conducted in a genotype blinded manner to mitigate potential bias. Image processing and data analysis was automated using imageJ macros. CA3 and cortex ROIs were drawn on DAPI channel while being blinded to other channels to reduce bias. ROI areas were compared between groups to ensure no differences.

Multiple controls were used for all key experiments to account for potential confounders. Raw *p-values* are reported for all comparisons up to 3-4 decimal points. All analysis is detailed in the materials and methods section to ensure reproducibility and transparency. A list of all the reagents used may be found in the supplementary tables.

## Supplementary figure and video legends

**Videos S1 and S2:** Regulatory volume decrease (RVD) response is reduced in Swell1 cKO microglia.

**Figure S1.**
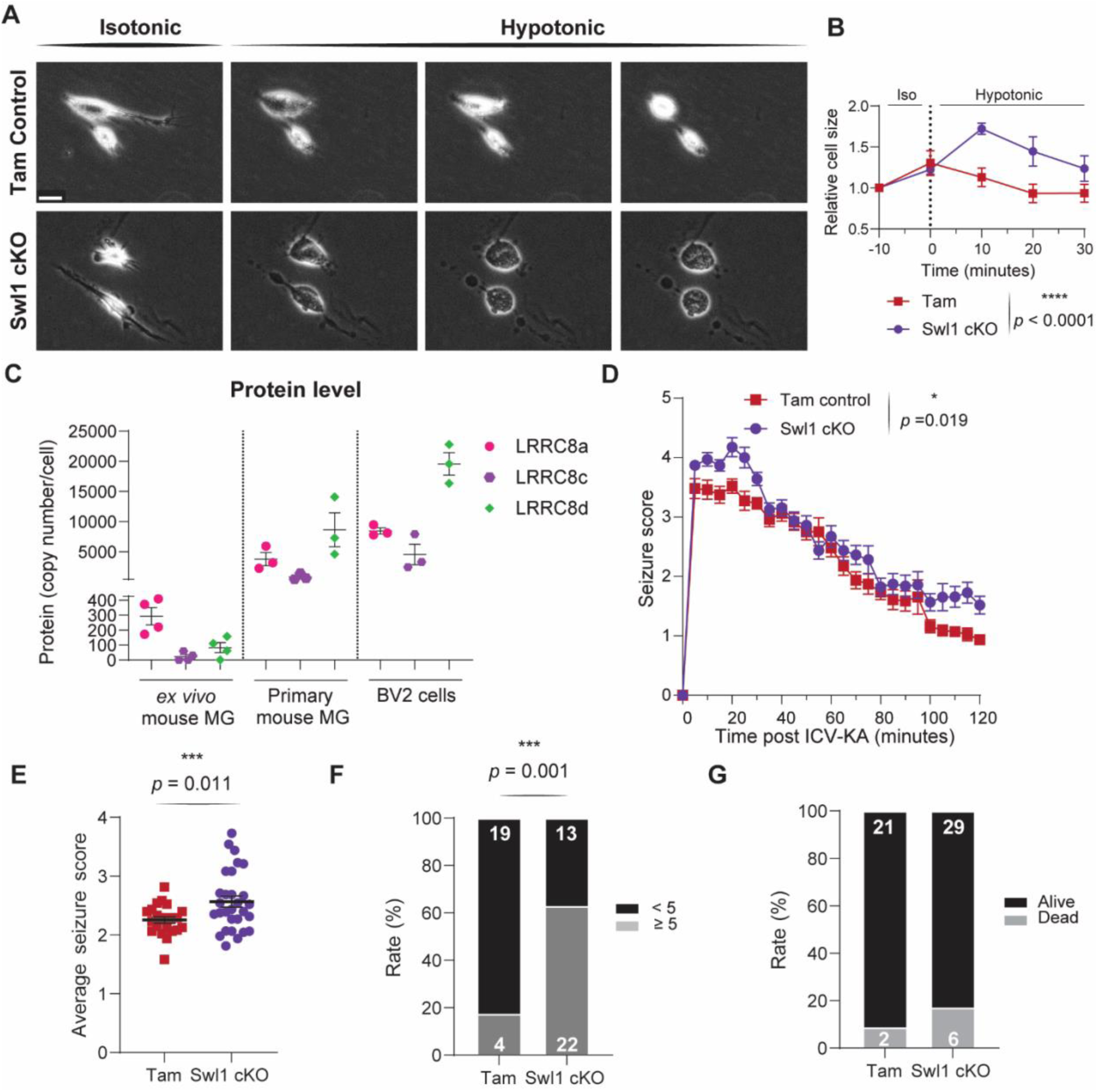
Regulatory volume decrease response is reduced in Swell1 cKO microglia. Swell1 cKO mice display worsened seizure compared to tamoxifen treated controls. A) Changes in microglia soma size and shape upon exposure to hypotonic medium. Scale bar, 20 µm. B) Quantification of changes in cell size (n=5 cells per group; statistic: 2-way ANOVA) C) Baseline protein levels for LRRC8a (Swell1), LRRC8c, and LRRC8d in *ex vivo* mouse microglia, cultured primary microglia, and BV2 cells (data from Lloyd et al., 2024). D) (D and E) Longitudinal and average seizure scores for Swell1 cKO vs tamoxifen control mice (n = 23-35 mice/ group; statistic: 2-way ANOVA and unpaired t-test) F) Percent of mice achieving a Racine score of 5 or higher at least once during the two-hour observation period (statistic: Chi-square test) G) Survival rate.

**Figure S2.**
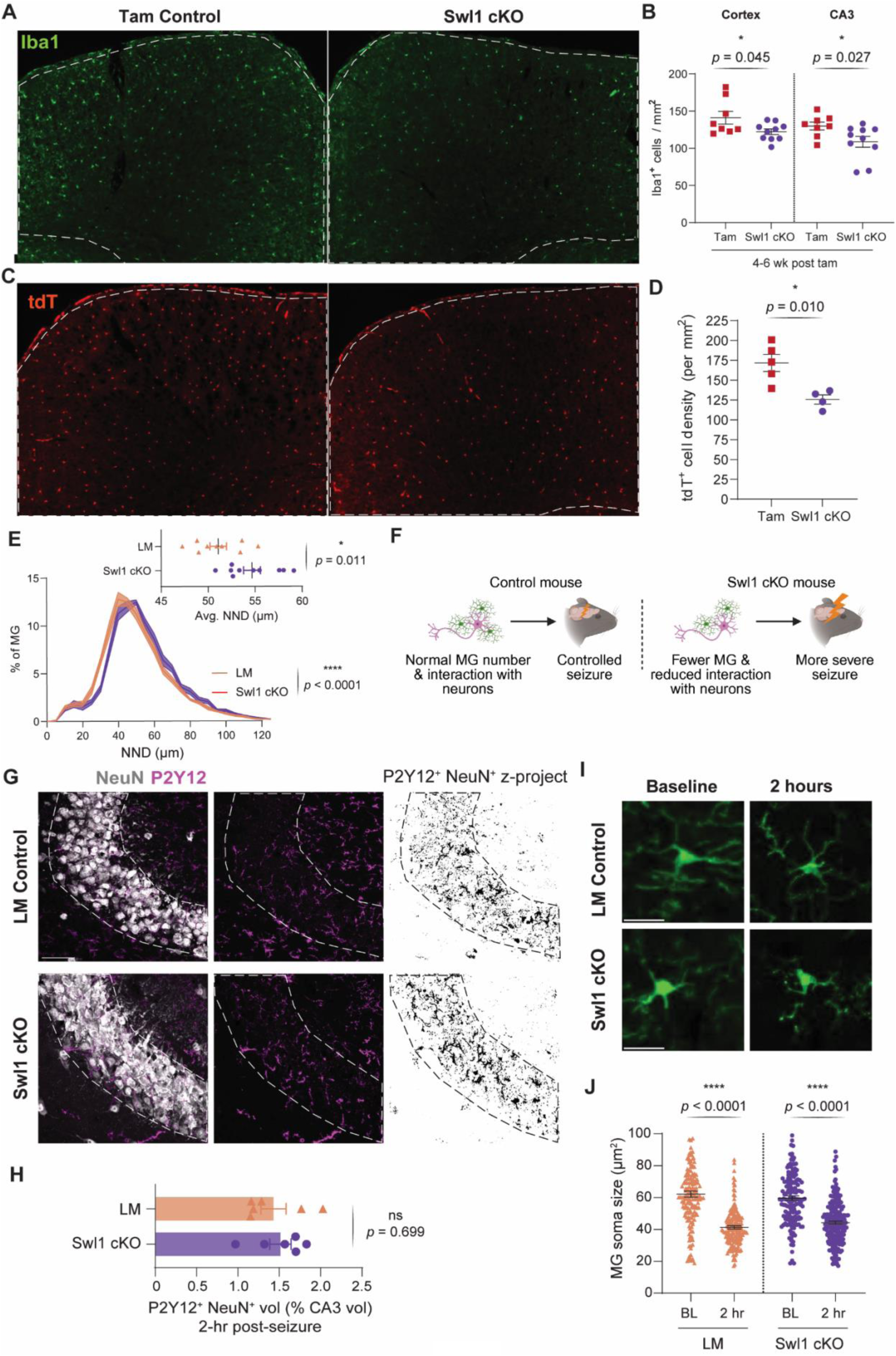
Microglial Swell1 cKO reduces microglia density but does not affect microglial-neuron physical interaction during acute seizure. A) Representative images of Iba1^+^ cells in the cortex. Scale bar, 50 µm. B) Quantification of Iba1 ^+^ cells in the cortex and CA3 (statistic: unpaired t-test and Mann-Whitney tests respectively) C) Representative images of tdTomato positive cells in the cortex. D) Quantification of tdT ^+^ cells in the cortex (statistic: unpaired t-test). E) Distribution and average of nearest neighbor distances for microglia (Iba1^+^ cells) in cortex. F) Microglia deficiency hypothesis of Swell1 mechanism in acute seizure G) Microglia (P2Y12)-neuron (NeuN) interaction in CA3 pyramidal layer at 2 hours after kainate induced seizure. Scale bar, 50 µm. H) P2Y12-NeuN double positive volume as a percent of CA3 pyramidal layer volume (statistic: Mann-Whitney test) I) (I and J) Representative images and quantification of changes in soma size at baseline and at 2-hours after seizure. Scale bar, 20 µm. (Statistic: nested t-test; dot, one microglia; 4-6 mice per group, 90-160 microglia per mouse)

**Figure S3.**
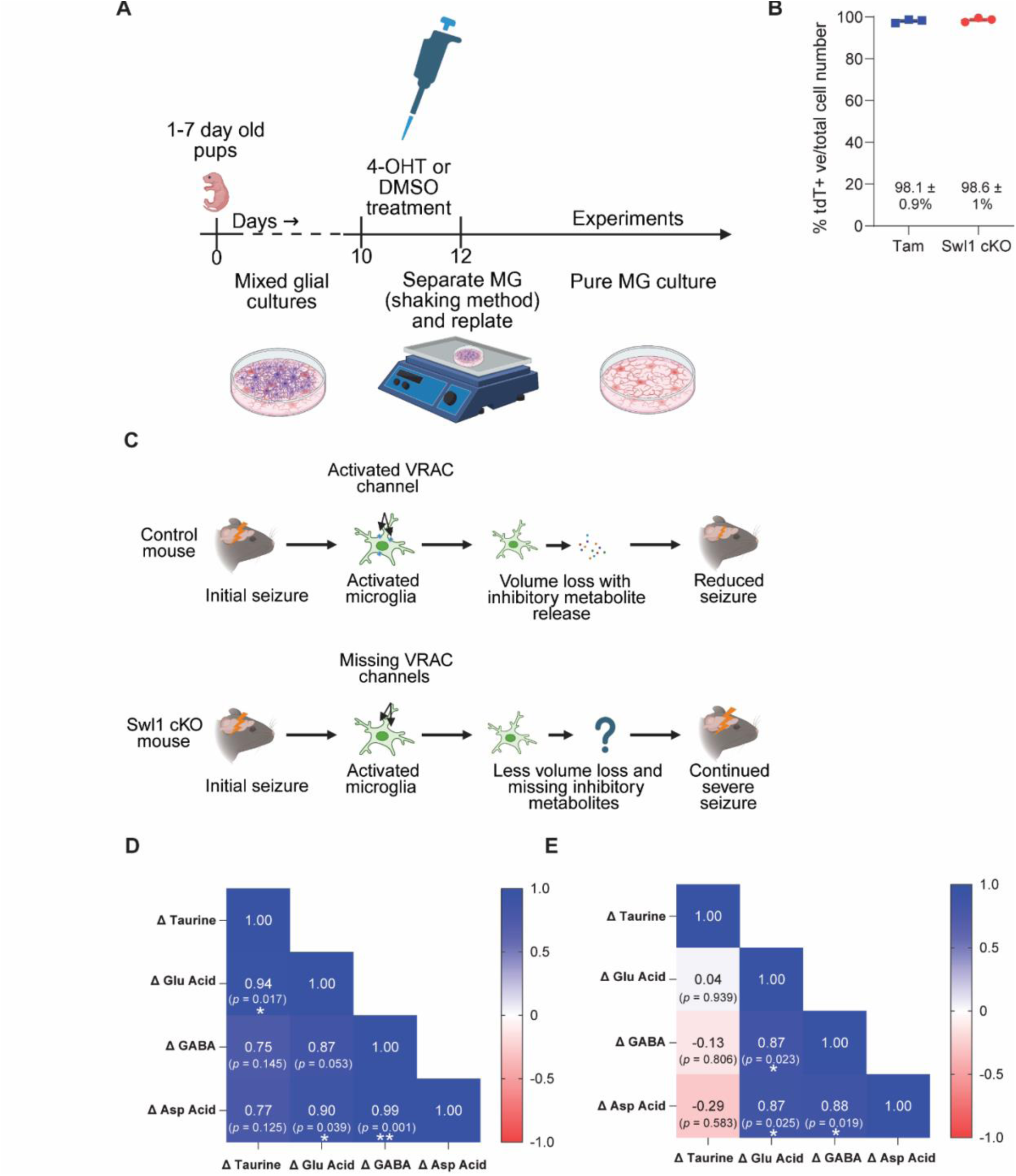
Primary microglia cultures are of high purity. Taurine dyshomeostasis is seen in post-seizure CSF of Swell1 cKO mice. A) Preparation of purified microglia cultures and 4-OHT treatment *in vitro* B) Culture purity assessed as % of cells positive for tdTomato signal C) Inhibitory neuromodulator hypothesis of Swell1 cKO mechanism in acute seizure D) (D and E) Correlated changes in CSF metabolites after seizure in littermate controls and Swell1 cKO mice respectively (delta values obtained by subtracting baseline average from post-seizure raw values; statistic: Pearson r).

**Figure S4.**
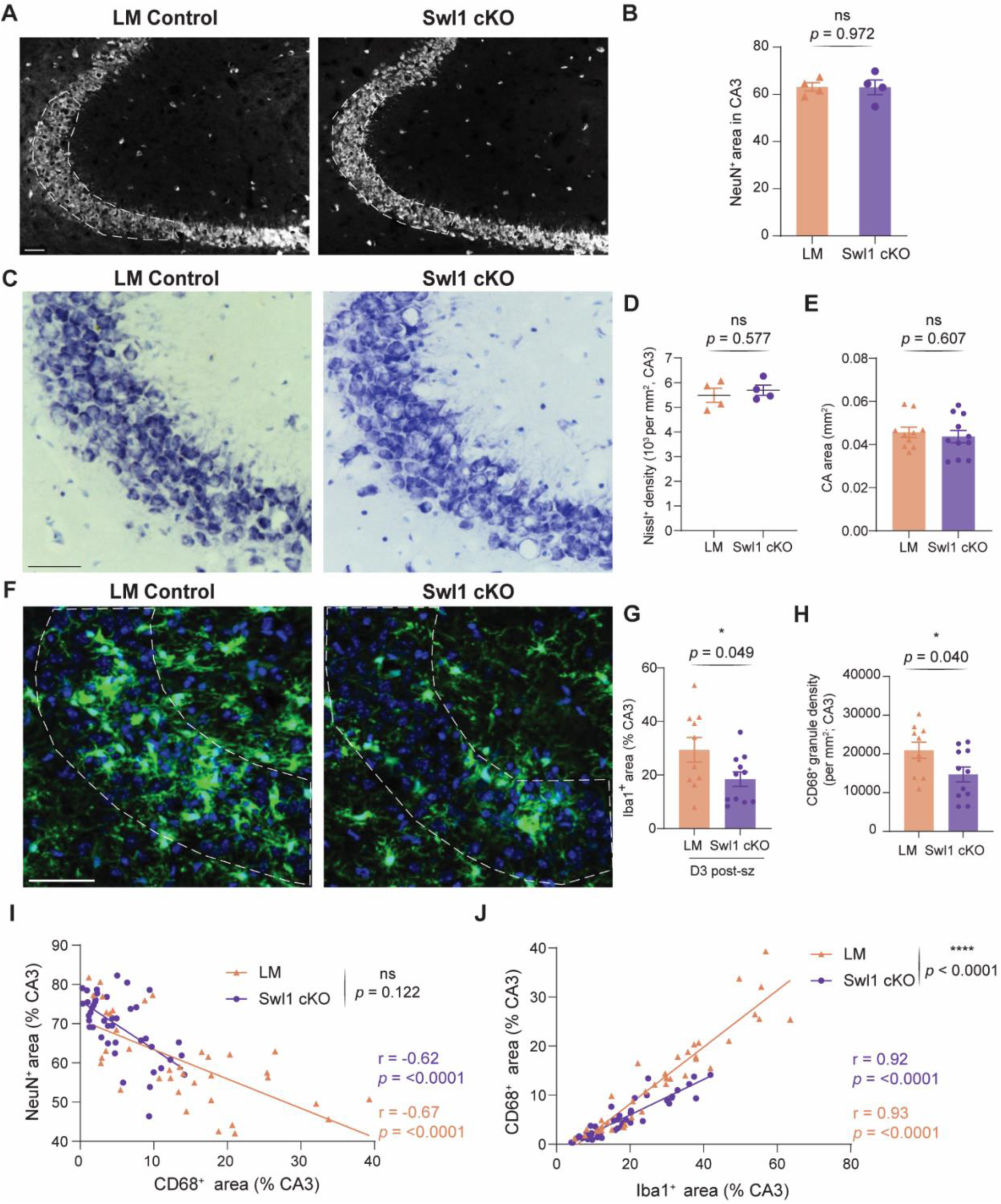
Baseline and post seizure (day 3) neuropathology in male mice. A) (A and B) Representative images and quantification of NeuN intensity at baseline (statistic: unpaired t-test). Scale bar, 50 µm. C) (C and D) Representative images and quantification of NeuN intensity at baseline (statistic: unpaired t-test). Scale bar, 50 µm. E) CA3 area measured at 3 days post-seizure (statistic: unpaired t-test). F) (F and G) Representative images and quantification of Iba1+ area in CA3 at day 3 post-seizure (statistic: unpaired t-test). Scale bar, 50 µm. H) Quantification of CD68^+^ granule density in CA3 pyramidal layer (statistic: unpaired t-test). I) Correlation between CD68 ^+^ and NeuN ^+^ area in CA3 pyramidal layer at day 3 after seizure (dot, one CA3 section; 3-5 sections examined per mouse; n = 10-11 mice/ group; statistic: statistic: Pearson correlation coefficient; simple linear regression to compare the slopes) J) Correlation between Iba1 ^+^ and CD68 ^+^ area in CA3 pyramidal layer at day 3 after seizure (dot, one CA3 section; 3-5 sections examined per mouse; n = 10-11 mice/ group; statistic: Pearson correlation coefficient; simple linear regression to compare the slopes) **Video S3 & S4**. Phagocytosis of opsonized beads is reduced in Swell1 cKO microglia.

**Figure S5.**
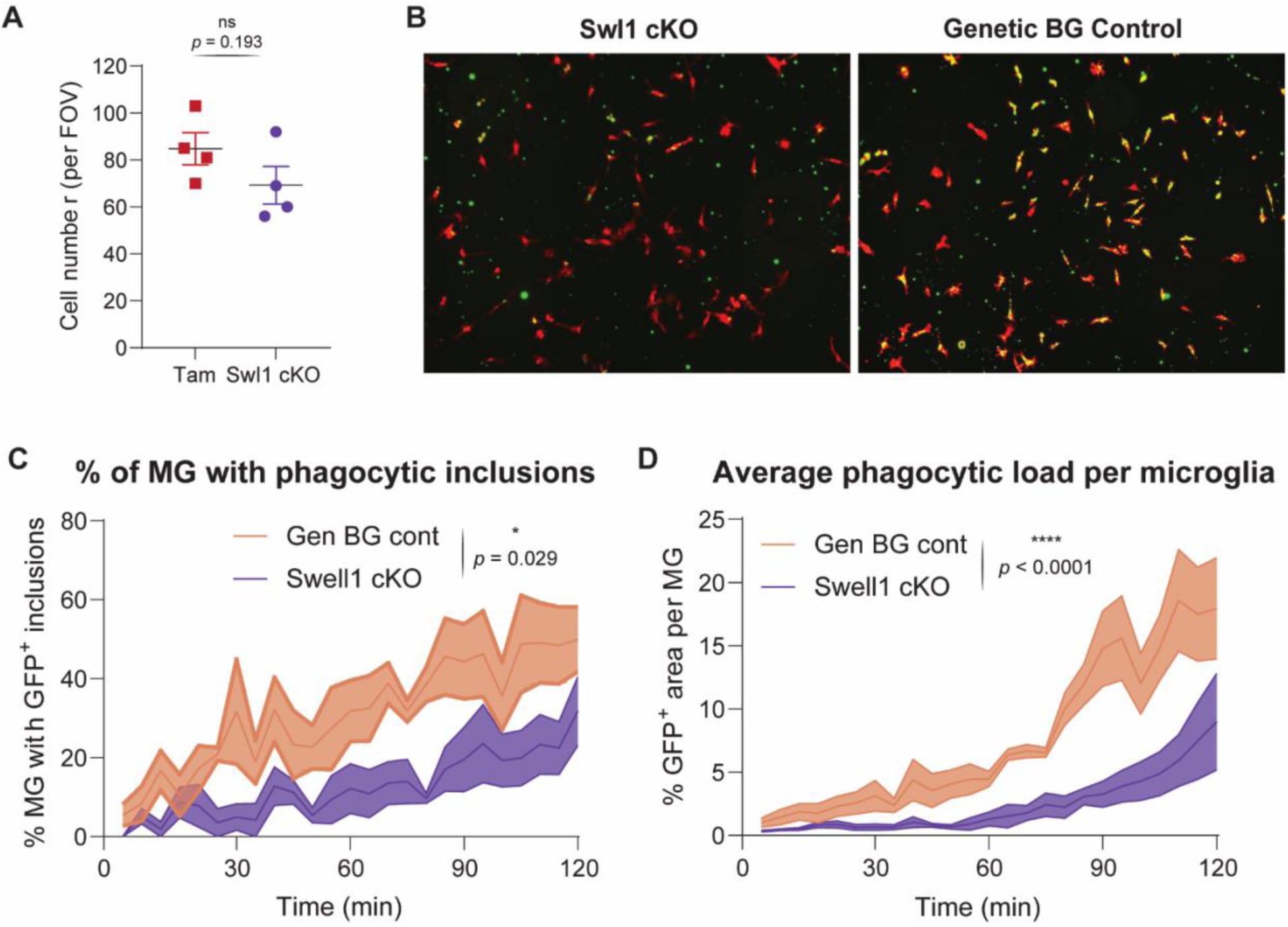
Phagocytic defects were confirmed in genotype matched cultured primary microglia. A) Cell counts at the time of bead phagocytosis assay shown in Fig. 5, D to F. B) Bead phagocytosis assay with genetic background matched controls. C) Quantification of percent microglia that are positive for phagocytic inclusions over a two-hour observation window (n = 3-4 wells/genotype; statistic: 2-way ANOVA) D) Quantification of average phagocytic load per microglia expressed as GFP^+^ area (n = 3-4 wells/genotype; results averaged per well; statistic: 2-way ANOVA)

**Figure S6.**
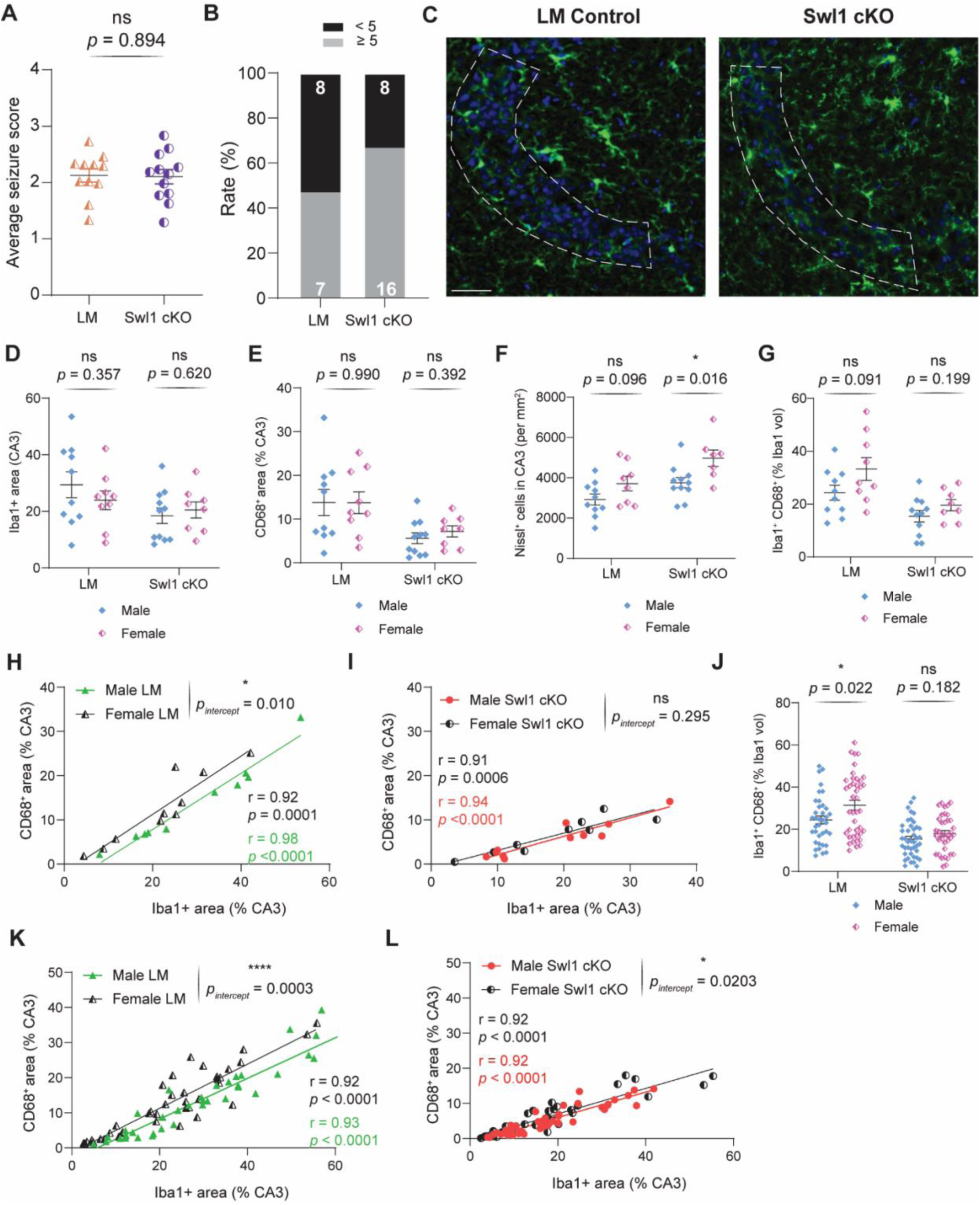
Female Swell cKO mice show no difference in seizure severity and a distinct neuropathology profile. A) Average Racine seizure scores over the two-hour observation period. (statistic: unpaired t-test) B) Percent of mice achieving a Racine score of 5 or higher at least once. C) Representative images of CA3 at baseline showing Iba1+ cells. Scale bar, 50 µm. D) (D to F) Iba1+ area, CD68+ area, and Nissl+ cell density in CA3 at day 3 post-seizure (dot, one animal; statistic: unpaired t-tests) G) Quantification of Iba1-CD68 double positive voxels in CA3 pyramidal layer as a percent of Iba1+ volume (dot, one animal; statistic: unpaired t-tests) H) (H and I) Sex wise correlations between Iba1^+^ and CD68 ^+^ areas in CA3 pyramidal layer at day 3 after seizure in LM and Swell1 cKO animals respectively (dot, one animal; statistic: Pearson correlation coefficient (r) and two-tailed *p*-value; simple linear regression to compare the slopes and intercept) J) (J to L) Same analysis as from G to I but with individual sections as data points (dot, one CA3 section; 3-5 sections examined per mouse; n = 9-10 mice/ group)

## Supplementary tables

**Table S1:** Metabolites values (in µM; mean ± SD) for primary microglia culture supernatants (n = number of wells)

|  |  | Tam controls |  | Swell1 cKO |  |
| --- | --- | --- | --- | --- | --- |
|  | Metabolite | Isotonic<br>(n = 5) | Hypotonic<br>(n = 5) | Isotonic<br>(n = 5) | Hypotonic<br>(n = 5) |
| 1 | Acetylcholine | <LLOD | <LLOD | <LLOD | <LLOD |
| 2 | Adenosine | <LLOD | 0.004 ± 0.001 | <LLOD | 0.004 ± 0.000 |
| 3 | Histidine | 0.033 ± 0.019 | 0.025 ± 0.009 | 0.050 ± 0.034 | 0.032 ± 0.010 |
| 4 | Serine | 0.114 ± 0.096 | 0.050 ± 0.049 | 0.214 ± 0.110 | 0.052 ± 0.045 |
| 5 | Taurine | 0.142 ± 0.043 | 1.543 ± 0.227 | 0.091 ± 0.021 | 0.716 ± 0.113 |
| 6 | Glutamine | 0.193 ± 0.075 | 0.141 ± 0.057 | 0.214 ± 0.094 | 0.130 ± 0.058 |
| 7 | Glycine | <LLOD | <LLOD | <LLOD | <LLOD |
| 8 | Aspartic Acid | 0.026 ± 0.018 | 0.057 ± 0.038 | 0.025 ± 0.027 | 0.028 ± 0.013 |
| 9 | Glutamic Acid | 0.078 ± 0.020 | 0.401 ± 0.117 | 0.063 ± 0.019 | 0.198 ± 0.026 |
| 10 | GABA | 0.004 ± 0.000 | 0.037 ± 0.008 | 0.005 ± 0.001 | 0.020 ± 0.002 |
| 11 | Norepinephrine | <LLOD | <LLOD | <LLOD | <LLOD |
| 12 | Dopamine | <LLOD | <LLOD | <LLOD | <LLOD |
| 13 | Epinephrine | 0.018 ± 0.005 | 0.026 ± 0.006 | 0.025 ± 0.011 | 0.023 ± 0.003 |
| 14 | Serotonin | <LLOD | <LLOD | <LLOD | <LLOD |

LLOD: Lower limit of detection
**Note:** If 50% or more values were below LLOD for an analyte in any group, they were discarded from analysis.

**Table S2:**
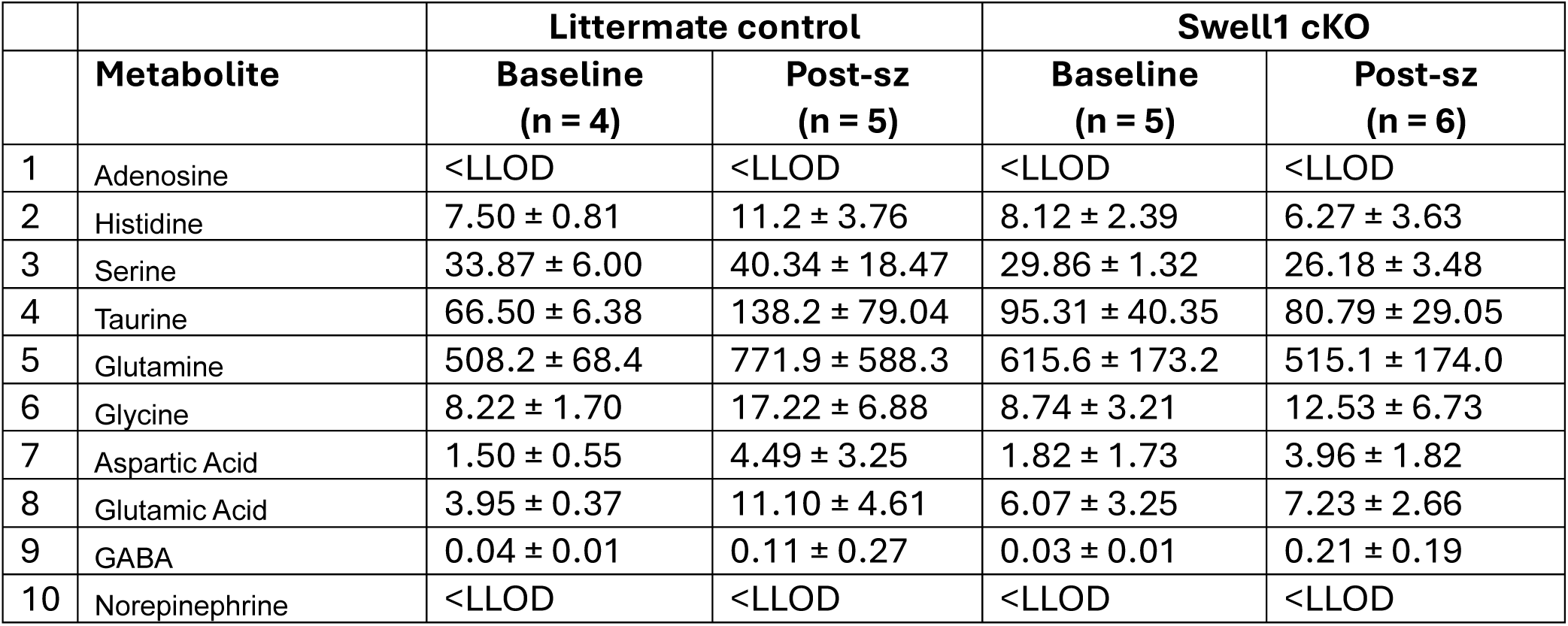
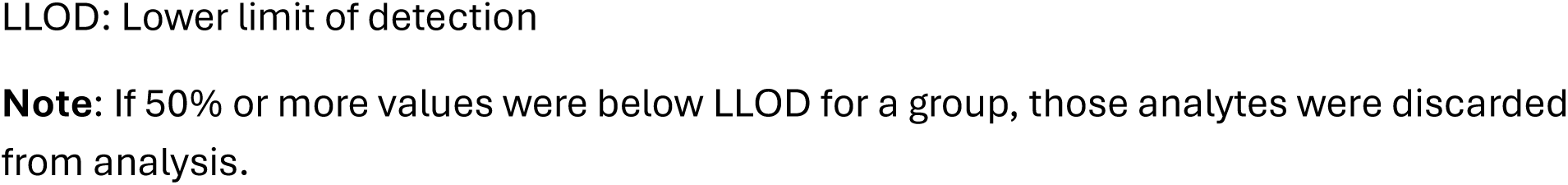
CSF metabolites values (in µM; mean ± SD) for male mice (n = number of mice)

|  |  | Littermate control |  | Swell1 cKO |  |
| --- | --- | --- | --- | --- | --- |
|  | Metabolite | Baseline<br>(n = 4) | Post-sz<br>(n = 5) | Baseline<br>(n = 5) | Post-sz<br>(n = 6) |
| 1 | Adenosine | <LLOD | <LLOD | <LLOD | <LLOD |
| 2 | Histidine | 7.50 ± 0.81 | 11.2 ± 3.76 | 8.12 ± 2.39 | 6.27 ± 3.63 |
| 3 | Serine | 33.87 ± 6.00 | 40.34 ± 18.47 | 29.86 ± 1.32 | 26.18 ± 3.48 |
| 4 | Taurine | 66.50 ± 6.38 | 138.2 ± 79.04 | 95.31 ± 40.35 | 80.79 ± 29.05 |
| 5 | Glutamine | 508.2 ± 68.4 | 771.9 ± 588.3 | 615.6 ± 173.2 | 515.1 ± 174.0 |
| 6 | Glycine | 8.22 ± 1.70 | 17.22 ± 6.88 | 8.74 ± 3.21 | 12.53 ± 6.73 |
| 7 | Aspartic Acid | 1.50 ± 0.55 | 4.49 ± 3.25 | 1.82 ± 1.73 | 3.96 ± 1.82 |
| 8 | Glutamic Acid | 3.95 ± 0.37 | 11.10 ± 4.61 | 6.07 ± 3.25 | 7.23 ± 2.66 |
| 9 | GABA | 0.04 ± 0.01 | 0.11 ± 0.27 | 0.03 ± 0.01 | 0.21 ± 0.19 |
| 10 | Norepinephrine | <LLOD | <LLOD | <LLOD | <LLOD |

LLOD: Lower limit of detection
**Note:** If 50% or more values were below LLOD for a group, those analytes were discarded from analysis.

**Table S3:** CSF metabolites values (8 in µM; mean ± SD) for female mice (n = number of mice)

|  | **Metabolite** | **Littermate control** |  | **Swell1 cKO** |  |
| --- | --- | --- | --- | --- | --- |
|  |  | **Baseline**<br>**(n = 4)** | **Post-sz**<br>**(n = 3)** | **Baseline**<br>**(n = 5)** | **Post-sz**<br>**(n = 2)** |
| 1 | Adenosine | 0.13 ± 0.12 | <LLOD | 0.07 ± 0.05 | 0.28 ± 0.21 |
| 2 | Histidine | 6.16 ± 6.81 | 5.68 ± 2.81 | 6.41 ± 3.77 | 10.69 ± 6.49 |
| 3 | Serine | 32.57 ± 30.94 | 20.75 ± 3.90 | 31.91 ± 19.33 | 48.63 ± 26.17 |
| 4 | Taurine | 72.19 ± 52.35 | 60.03 ± 42.90 | 82.40 ± 22.49 | 115.8 ± 90.8 |
| 5 | Glutamine | 551.9 ± 538.0 | 421.7 ± 93.6 | 617.3 ± 430.6 | 961.6 ± 511.8 |
| 6 | Glycine | 5.28 ± 1.86 | 8.78 ± 4.70 | 8.35 ± 4.36 | 5.90 ± 3.69 |
| 7 | Aspartic Acid | 2.38 ± 0.59 | 3.45 ± 1.59 | 2.33 ± 1.11 | 2.40 ± 1.35 |
| 8 | Glutamic Acid | 4.25 ± 0.75 | 6.73 ± 3.56 | 4.28 ± 1.37 | 4.40 ± 2.62 |
| 9 | GABA | 0.15 ± 0.03 | 0.17 ± 0.13 | 0.08 ± 0.04 | 0.09 ± 0.03 |
| 10 | Norepinephrine | <LLOD | <LLOD | <LLOD | <LLOD |

**Table S4:** Plasma metabolites values (in µM; mean ± SD) for male mice (n = number of mice)

|  | **Metabolite** | **Littermate control** |  | **Swell1 cKO** |  |
| --- | --- | --- | --- | --- | --- |
|  |  | **Baseline**<br>**(n = 4)** | **Post-sz**<br>**(n = 6)** | **Baseline**<br>**(n = 5)** | **Post-sz**<br>**(n = 6)** |
| 1 | Adenosine | 0.24 ± 0.15 | 0.13 ± 0.11 | 0.13 ± 0.08 | 0.11 ± 0.06 |
| 2 | Histidine | 116.8 ± 50.4 | 127.2 ± 29.3 | 103.9 ± 18.2 | 107.6 ± 17.27 |
| 3 | Serine | 205.8 ± 68.3 | 200.5 ± 36.3 | 179.8 ± 24.29 | 169.7 ± 57.0 |
| 4 | Taurine | 836.8 ± 304.5 | 891.7 ± 376.6 | 639.2 ± 114.5 | 552.7 ± 182.5 |
| 5 | Glutamine | 946.4 ± 444.5 | 870.2 ± 187.7 | 775.8 ± 66.2 | 689.2 ± 134.4 |
| 6 | Glycine | 490.2 ± 84.5 | 453.0 ± 112.9 | 435.5 ± 79.6 | 295.5 ± 74.4 |
| 7 | Aspartic Acid | 6.83 ± 4.84 | 10.19 ± 5.94 | 3.71 ± 1.59 | 7.84 ± 3.99 |
| 8 | Glutamic Acid | 77.55 ± 30.16 | 106.6 ± 92.9 | 31.94 ± 10.38 | 35.11 ± 22.54 |
| 9 | GABA | 0.55 ± 0.17 | 1.14 ± 1.10 | 0.27 ± 0.06 | 0.52 ± 0.75 |
| 10 | Norepinephrine | 0.10 ± 0.01 | 0.12 ± 0.07 | 0.10 ± 0.01 | 0.10 ± 0.06 |

**Table S5:** Plasma metabolites values (in µM; mean ± SD) for female mice (n = number of mice)

|  |  | Littermate control |  | Swell1 cKO |  |
| --- | --- | --- | --- | --- | --- |
|  | Metabolite | Baseline<br>(n = 6) | Post-sz<br>(n = 3) | Baseline<br>(n = 6) | Post-sz<br>(n = 2) |
| 1 | Adenosine | 0.11 ± 0.02 | 0.23 ± 0.05 | 0.11 ± 0.02 | 0.13 ± 0.01 |
| 2 | Histidine | 124.1 ± 16.3 | 117.6 ± 35.5 | 128.7 ± 24.1 | 151.7 ± 73.7 |
| 3 | Serine | 225.2 ± 34.4 | 207.7 ± 39.6 | 201.3 ± 27.3 | 300.0 ± 154.8 |
| 4 | Taurine | 781.4 ± 179.9 | 853.1 ± 125.0 | 757.3 ± 203.7 | 915.5 ± 202.5 |
| 5 | Glutamine | 889.8 ± 91.0 | 866.1 ± 251.8 | 849.1 ± 154.1 | 888.9 ± 328.7 |
| 6 | Glycine | 341.3 ± 81.54 | 428.0 ± 18.3 | 313.7 ± 65.5 | 455.2 ± 235.0 |
| 7 | Aspartic Acid | 13.20 ± 4.19 | 26.16 ± 12.76 | 10.44 ± 5.55 | 25.33 ± 9.56 |
| 8 | Glutamic Acid | 38.30 ± 10.38 | 78.92 ± 8.17 | 28.70 ± 13.84 | 52.71 ± 5.69 |
| 9 | GABA | 0.15 ± 0.05 | 0.50 ± 0.27 | 0.38 ± 0.36 | 0.19 ± 0.04 |
| 10 | Norepinephrine | 0.07 ± 0.01 | 0.17 ± 0.10 | 0.06 ± 0.01 | 0.07 ± 0.01 |

**Table S6:**
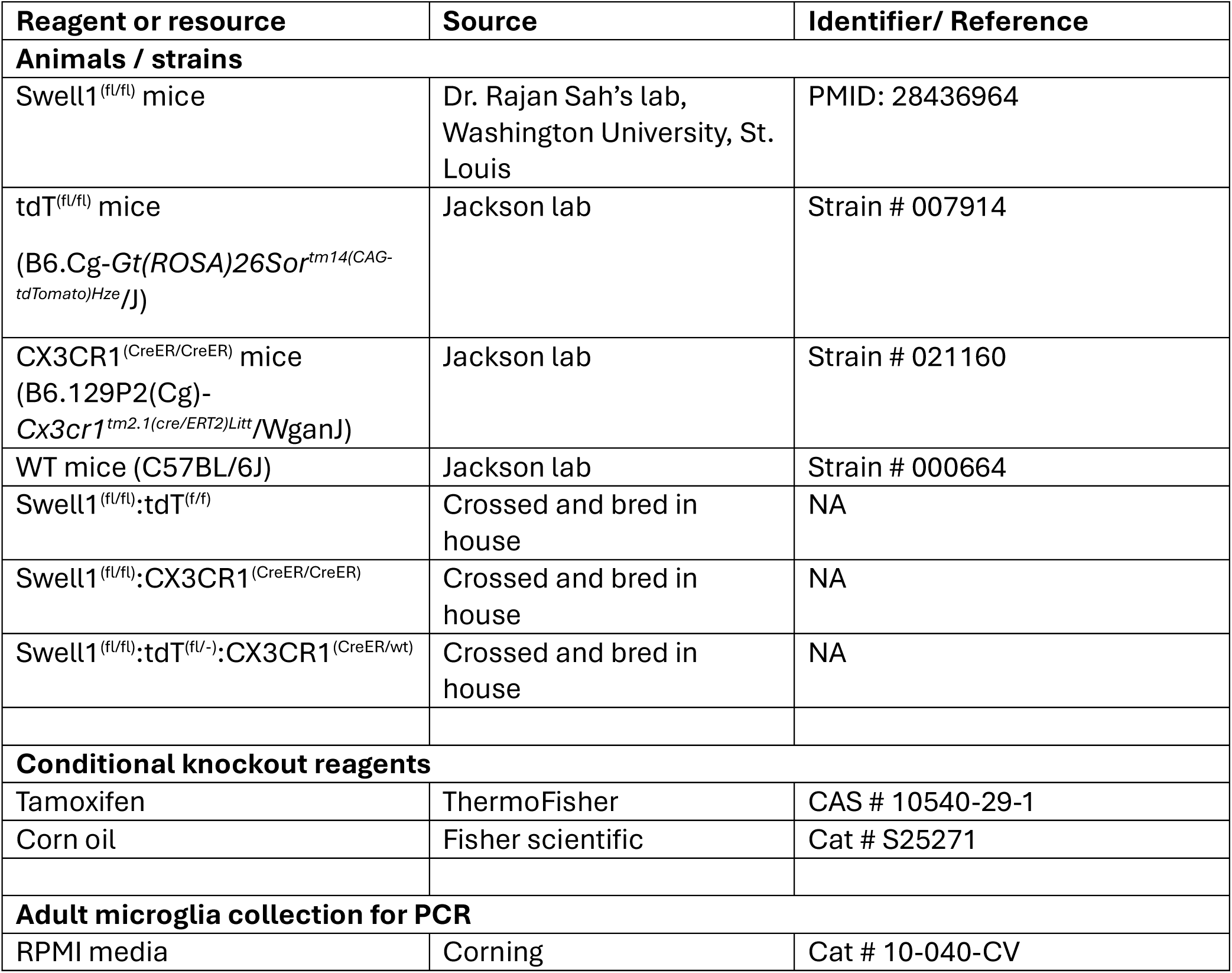
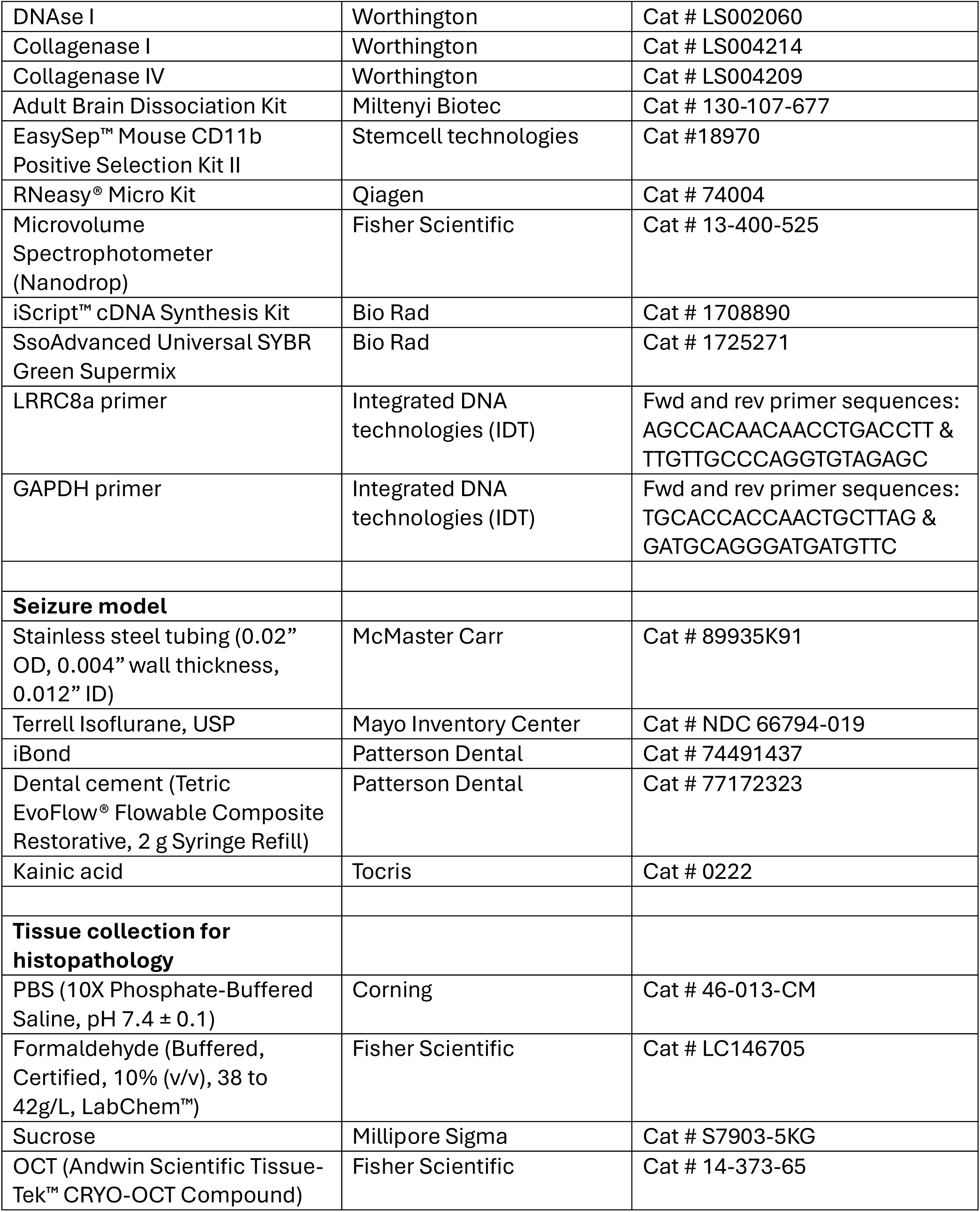
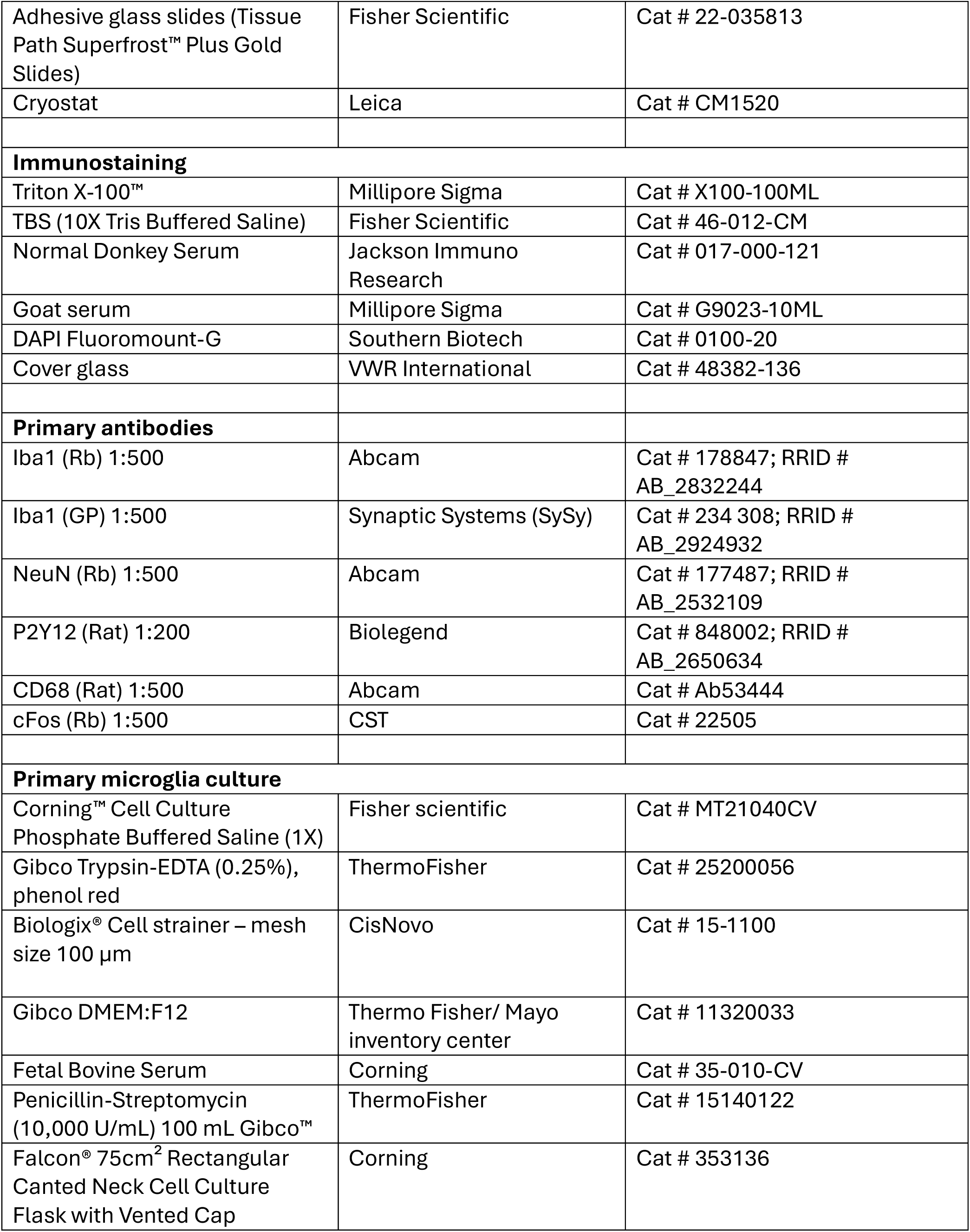
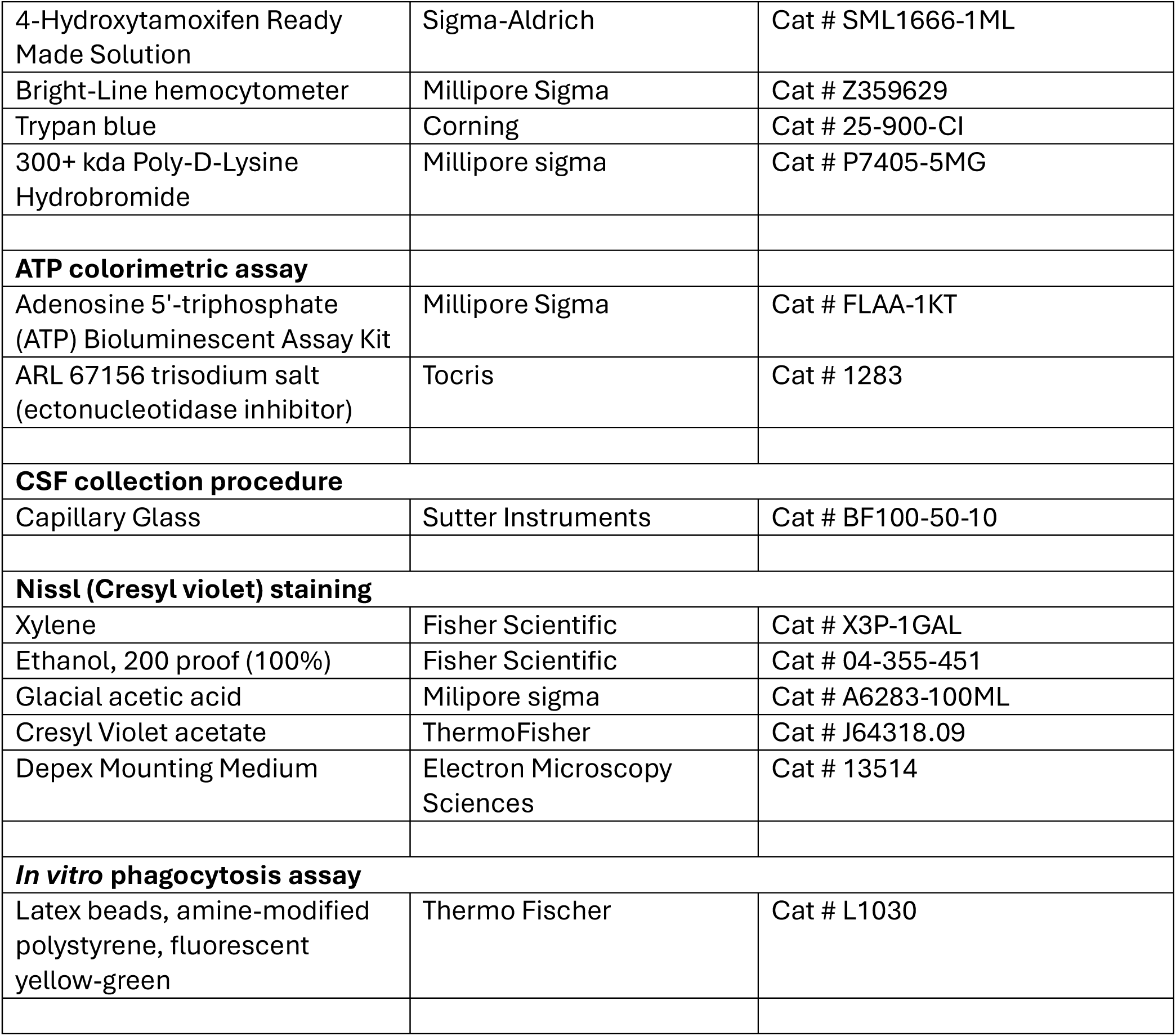
Resource table.

